# The Rab5 effector Rabankyrin-5 mediates endosomal fusion and trafficking of Human Papillomavirus during early entry

**DOI:** 10.1101/2025.08.05.668525

**Authors:** Madison Love, Jian Xie, Pengwei Zhang

## Abstract

The fusion of newly formed early endosomal vesicles after endocytosis is a crucial step in viral infection. It facilitates the transition of many viruses from viral internalization to downstream intracellular trafficking within the endosomal network, ultimately enabling their delivery to intracellular replication sites. Despite its significance, the molecular mechanisms regulating the fusion of these vesicles remain poorly understood. In this study, we show that Rabankyrin-5, a Rab5 effector, is essential for the fusion of human papillomavirus (HPV)-containing early endosomes during viral entry. Additionally, Rabankyrin-5 acts as a dynein adaptor, directly binding both the HPV minor capsid protein L2 and the dynein motor complex to link virus-carrying early endosomes to the dynein transport machinery, thereby promoting virus movement along microtubules. These dual functions enable the coordinated integration of endosomal fusion with microtubule-based transport during the early stages of viral entry.

## Introduction

Following the delivery of internalized cargo by endocytic carriers, fusion to form early endosomes is essential for endosomal sorting and maturation (1, 2). These processes are sequential and functionally linked within the endocytic pathway, working together to ensure that cargo is properly trafficked, sorted, and processed within the cell (2). For many viruses, successful entry similarly depends on the fusion of newly formed, virus-enriched endocytic vesicles with early endosomes, enabling subsequent endosomal escape and access to specific transport routes leading to sites of genome replication (3–6).

Human papillomaviruses (HPVs) enter host cells via endocytosis and are subsequently incorporated into the endosomal network (7, 8). Nascent HPV-containing endocytic carriers are transported to early endosomes and then traverse endosomal compartments, where the virus undergoes disassembly within late endosomes and is sorted into the retrograde transport pathway toward the Golgi apparatus (7). However, the mechanisms regulating the cytoplasmic transport of these virus-carrying vesicles for delivery to and fusion with early endosomes remain poorly understood.

HPV infection is responsible for nearly all cases of cervical cancer, as well as a significant proportion of oropharyngeal and anogenital cancers (9). The virus comprises a circular double-stranded DNA genome encased within a capsid formed by the major protein L1 and the minor protein L2. Following endocytosis, HPV remains within vesicular compartments as it traffics from the plasma membrane to the nucleus, where genome replication occurs (10–12). A key feature of L2 is a C-terminal cell-penetrating peptide (CPP) that facilitates membrane translocation, allowing portions of L2 to access the cytoplasm and interact with host cellular machinery (13, 14). Rab proteins and their effectors are among the host factors hijacked by HPV. During entry, L2 binds the endosomal sorting complex retromer and later dissociates from it in a process mediated by the cycling of the small GTPase Rab7, which directs the virus into the retrograde transport pathway from the endosome to the Golgi (15–17).

Rab proteins, a family of small GTPases, and their effectors play crucial roles in defining membrane identity and regulating membrane cargo transport (18, 19). Nascent endocytic vesicles become enriched in Rab5, which cooperates with its effectors to facilitate membrane targeting, tethering, and fusion between early endosomes, thereby promoting the formation and maturation of late endosomes (20–22). Previous studies have shown that dominant-negative (DN) and constitutively active (CA) mutants of Rab5A recombinant proteins lead to reduced HPV infection (23, 24). However, the precise mechanisms by which Rab5 mediates the early stages of HPV entry remain unknown. Rabankyrin-5, a Rab5 effector, was identified as a high hit in a genetic screen for human genes essential in HPV16 infection (25). It interacts with Rab5 to regulate retromer-mediated retrograde transport of cellular cargos, but whether it contributes to endosomal fusion remains unclear (26–29).

A recently proposed model of endosomal maturation suggests that distinct very early endosomal populations undergo fusion and maturation at different rates, depending on their motility (30–32). This idea is supported by studies on epidermal growth factor receptor (EGFR) trafficking, which depends on the microtubule motor protein dynein for proper localization to fuse and mature endosomes (30, 33). Dynein is a multi-subunit motor protein that converts chemical energy from ATP hydrolysis into mechanical force. In coordination with its cofactor dynactin, it moves along microtubules to transport a wide variety of cargoes. Cargo specificity and functional diversity are mediated by dynein adaptor proteins, which link dynein to distinct cargo types (19). Among these, certain Rab effectors function as adaptors by bridging the motor complex to Rab-decorated membranes. This interaction promotes dynein-driven motility and enables efficient transport of membrane-bound cargo along microtubules (19).

HPV exploits the host microtubule network during infection by utilizing dynein to travel through the cytoplasm toward the nucleus (34–36). Given that microtubules not only provide tracks for the movement of cargo-enriched early endosomes but also are involved in fusion processes critical for endosome maturation (19, 21, 33), it is plausible that the dynein transport machinery contributes to both the trafficking and fusion of virus-containing early endosomes during the early stages of HPV entry. In this study, we found that Rabankyrin-5 is essential for the fusion of virus-carrying early endosomes and acts as a previously unrecognized dynein adaptor. It mediates the physical connection of dynein to HPV-carrying early endosomes through direct interactions with both the motor complex and the L2 protein. These interactions promote the directed motility of early endosomes and enhance their fusion capacity. Our findings reveal that endosomal motility is not merely a consequence of transport but is required to coordinate membrane fusion events that are critical for productive HPV entry. This work provides mechanistic insights into how endocytic trafficking and membrane fusion are functionally integrated during the early stages of viral infection.

## Results

### Rabankyrin-5 is required for HPV retrograde transport and is associated with HPV

Rab5 and its effectors, enriched on nascent early endosomes, form a key complex that regulates endosomal trafficking and fusion, which are essential for endosome formation and maturation (1, 2). Since both DN and CA Rab5A mutants reduce HPV infection (23), we examined HPV infectivity in Rab5A-depleted cells. Three human cell types were used: HeLa, HaCaT, and human foreskin keratinocytes (HFKs) immortalized via retroviral transduction of the E6 and E7 oncogenes (13, 37). HPV infection was measured using a well-established pseudovirus (PsV) system consisting of the L1 and FLAG-tagged L2 capsid proteins of HPV16, encapsulating a reporter plasmid encoding HcRed, thereby allowing quantitative measurement of infection efficiency (38). Cells were transfected with either non-targeting control siRNA or Rab5A-specific siRNA. Rab5A knockdown resulted in a significant reduction in HPV infection across all three cell types (Fig. 1A). Knockdown efficiency was confirmed by quantitative real-time PCR of Rab5A mRNA levels (Fig. S1A). Given that Rabankyrin-5, a Rab5 effector, was identified as a hit in a genetic screen for host factors essential for HPV16 infection (25), we next evaluated its role in the infection process. Similar to Rab5A, the depletion of Rabankyrin-5 led to a significant reduction in HPV infection in all three cell types (Fig. 1B and Fig. S1B).

**Figure 1.**
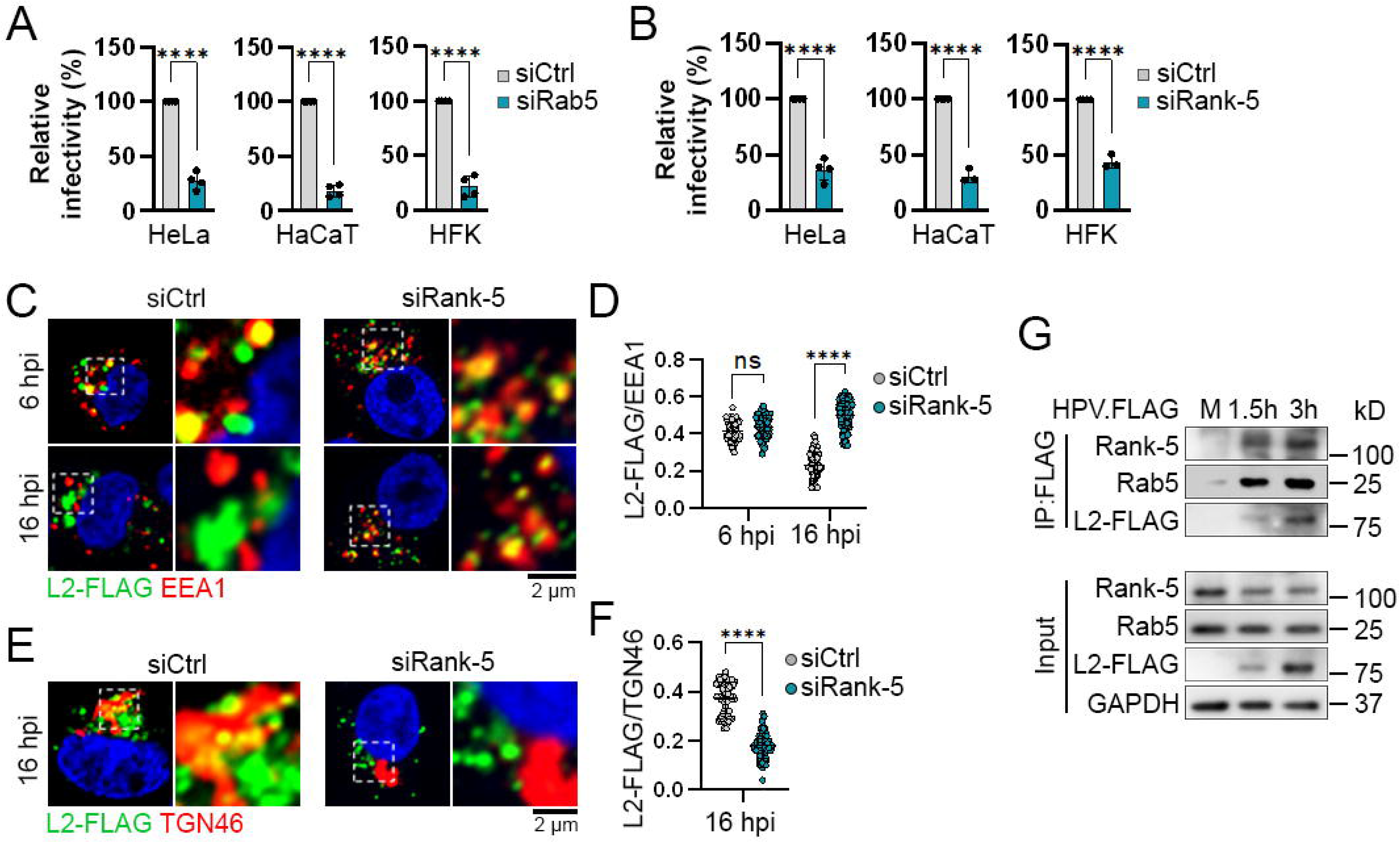
Rabankyrin-5 is required for HPV retrograde transport and is associated with HPV. (A and B) HeLa, HaCaT, and HFK cells were transfected with negative control siRNA (siCtrl) or siRNA targeting Rab5A (A) or Rabankyrin-5 (Rank-5) (B) for 48 hours. The cells were then mock-infected or infected at an MOI of 2 with HPV.FLAG PsVs containing the HcRed reporter plasmid. At 48 hpi, HcRed fluorescence was measured by flow cytometry. Results are presented as percent relative infectivity, based on the mean of fluorescence intensity, normalized to siCtrl-treated cells (set at 100%). Bars and error bars represent the mean and standard deviation (SD), respectively. Statistical significance was assessed using a two-tailed *t*-test with unequal variance, based on three independent experiments, in comparison to siCtrl-treated cells. ****p < 0.0001. (C-F) HeLa cells were transfected with siCtrl or siRNA targeting Rabankyrin-5. After 48 hours, cells were mock-infected or infected with HPV.FLAG PsVs at an MOI of 25. At 6 or 16 hpi, cells were stained with anti-FLAG, anti-EEA1 (C and D), or anti-TGN46 (E and F) antibodies. Fluorescence was visualized by confocal microscopy. (C and E) Representative images (left) and insets (right) are shown. L2-FLAG is shown in green; EEA1, in red (C); TGN46, in red (E); and nuclei, in blue. Co-localization of L2-FLAG with EEA1 (C) or TGN46 (E) is pseudocolored in yellow. Inset scale bar: 2 μm. (D and F) The co-localization signal was quantified using Fiji software, based on the Pearson’s correlation coefficient, from at least 60 cells in three independent experiments. Results are presented as mean ± SD. Each dot represents an individual cell. ****p < 0.0001; ns, not significant. (G) HeLa cells were mock-infected (M) or infected with HPV.FLAG at an MOI of 25 for the indicated time points. Cell lysates were subjected to immunoprecipitation using M2 FLAG beads, and the immunoprecipitated samples were analyzed by immunoblotting with anti-FLAG, anti-Rab5, and anti-Rabankyrin-5 antibodies.

To determine the specific step of HPV entry affected by Rabankyrin-5 knockdown, immunofluorescence analysis was performed in knockdown cells to track viral trafficking by examining the colocalization of the virus with cellular organelle markers. As shown in Fig. 1C and D, in control cells, L2-FLAG clearly colocalized with the early endosome marker EEA1 at 6 hours post-infection (hpi). By 16 hpi, this colocalization with EEA1 decreased, while colocalization with the trans-Golgi network (TGN) marker TGN46 increased (Fig. 1E and F).

This pattern reflects the retrograde transport of incoming viruses from endosomes to the TGN. In contrast, Rabankyrin-5-depleted cells exhibited significant colocalization of L2 with EEA1 but not with TGN46 at 16 hpi, suggesting retention of the virus in early endosomes (Fig. 1C-F). This accumulation was further confirmed using a proximity ligation assay (PLA), which detects proteins in close proximity (within ∼40 nm) by generating a fluorescent signal at the single-molecule level. HeLa cells were infected with HPV16 PsVs for 6 and 16 hours and subjected to PLA to assess the proximity of L1 with either EEA1 or TGN46 (Fig. S1C and D). In control siRNA-treated cells, PLA signals for EEA1 and L1 decreased at 16 hpi, while signals for TGN46 and L1 increased, consistent with the trafficking of viral particles from endosomes to the TGN. In contrast, Rabankyrin-5 knockdown resulted in an increase, rather than a decrease, in the PLA signal between EEA1 and L1 at 16 hpi, confirming the retention of viruses in the early endosomes.

We next determined whether Rabankyrin-5 physically associates with HPV during entry using co-immunoprecipitation (co-IP) assays. HeLa cells were either mock-infected or infected with FLAG-tagged HPV16 PsVs (HPV.FLAG). At 1.5 and 3 hpi, cell lysates were immunoprecipitated using anti-FLAG affinity gel. As shown in Fig. 1G, both Rab5 and Rabankyrin-5 co-precipitated with L2-FLAG at these time points. Since endocytic vesicles develop into early endosomes as Rab5 accumulates on their surface, these findings suggest that Rabankyrin-5 associates with HPV at the early endosome and plays an essential role in a step preceding viral escape from the endosome.

### Rabanykrin-5 mediates HPV-carrying early endosomal fusion

After internalization, endocytic vesicles convert into early endosomes and increase in size through fusion, forming larger early endosomal structures (1, 22). Interestingly, in cells depleted of Rabankyrin-5, we observed smaller HPV L2 puncta at both 6 and 16 hpi (Fig. 1C, 1E, 2A, and Fig. S2A). To investigate this further, we measured the size of HPV L2 puncta in Rabankyrin-5-depleted HeLa cells (Fig. 2B) and HaCaT cells (Fig. S2B) at 6 hpi. Quantitative analysis revealed that a large population of L2 puncta in Rabankyrin-5-depleted cells was approximately 2-3 times smaller (0.15 μm^2^ in HeLa cells and 0.32 μm^2^ in HaCaT cells) than those in control cells (0.42 μm^2^ in HeLa cells and 0.87 μm^2^ in HaCaT cells) (Fig. 2B and C; Fig. S2B and C). As EEA1-positive early endosomes in Rabankyrin-5-depleted cells exhibited similar morphology to those in control cells (Fig. 1C), Rabankyrin-5 depletion may not impair early endosome integrity but rather disrupts the fusion of virus-containing early endosomes during endosome formation or maturation.

**Figure 2.**
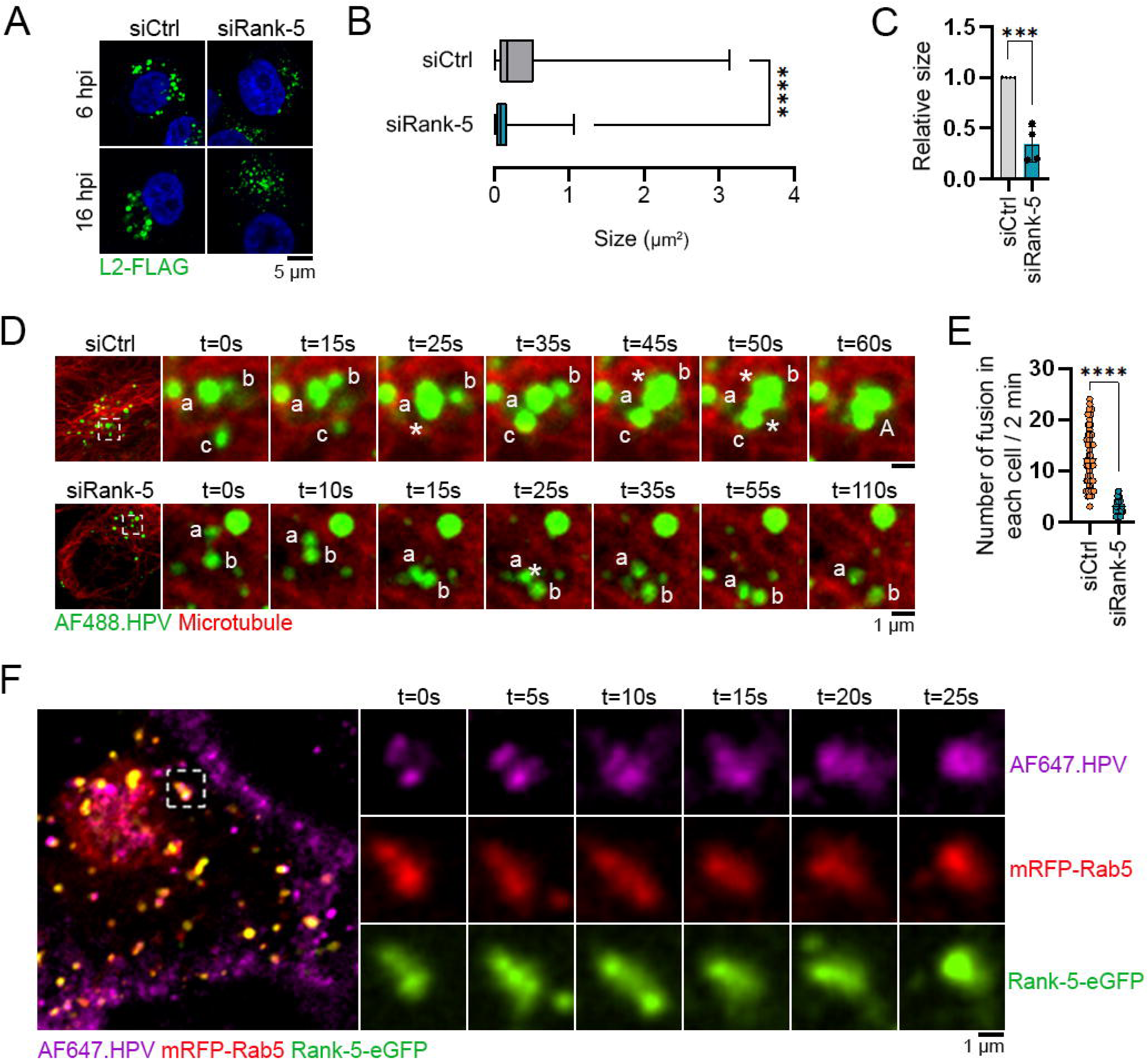
Rabankyrin-5 contributes to the fusion of HPV-carrying early endosomes. (A) HeLa cells were transfected with negative control siRNA (siCtrl) or siRNA targeting Rabankyrin-5 (Rank-5). After 48 hours, cells were mock-infected or infected with HPV.FLAG PsVs at an MOI of 25. At 6 and 16 hpi, cells were stained with an anti-FLAG antibody, and fluorescence was visualized by confocal microscopy. Representative images are shown. L2-FLAG is shown in green; nuclei are shown in blue. Scale bar: 5 μm. (B) The size of virus particles in infected cells at 6 hpi was analyzed and plotted from three independent experiments. Bars indicate the full range of values. ****p < 0.0001; n = 1000-1200 particles. (C) The relative size was determined based on the average size of virus particles shown in (B) from four independent experiments, normalized to siCtrl-treated cells (set at 1). Each dot represents the mean particle size from a single experiment. Statistical significance was assessed using a two-tailed *t*-test with unequal variance, comparing each condition to siCtrl-treated cells. ***p < 0.001. (D) HeLa cells were transfected with siRNAs as described in (A) and infected with AlexaFluor 488-labeled HPV.FLAG PsVs (AF488.HPV, green) at an MOI of 100. At 1.5 hpi, microtubules were visualized by staining prior to image acquisition. Representative time-lapse insets are shown on the right. Lowercase letters indicate distinct virus particles; asterisks denote collisions and fusion events; uppercase letters indicate larger fused particles. Microtubules are pseudocolored in red. Inset scale bar: 1 μm. (E) The number of fusion events per cell over a 2-minute interval, as observed in panel (D), was quantified from three independent experiments and plotted. Each dot represents an individual cell. n = 50-60 cells; ****p < 0.0001. (F) HeLa cells were transfected with mRFP-Rab5 (red) and Rank-5-eGFP (green). After 24 hours, cells were infected with AlexaFluor 647-labeled HPV.FLAG PsVs (AF647.HPV, magenta). Representative time-lapse insets are shown on the right. Inset scale bar: 1 μm.

The fusion process includes vesicle capture and tethering by the target membrane, followed by SNARE priming, membrane docking, and membrane fusion (22, 29, 39). Rab GTPases and their effectors function as targeting and tethering proteins that link two vesicular transport carriers together prior to membrane fusion (21, 22, 29). To assess the role of Rabankyrin-5 in the fusion of virus-carrying vesicles, we employed live-cell imaging in HeLa cells using fluorophore-labeled HPV16 PsVs (40). The purified HPV.FLAG PsVs were covalently labeled with the fluorophores AlexaFluor488 or AlexaFluor647. The labeled PsVs exhibited comparable purity, contained similar levels of L1 and L2 proteins, and showed infectivity equivalent to that of unlabeled HPV16 PsVs (Fig. S2D and E). HeLa cells were treated with either control siRNA or Rabankyrin-5-targeting siRNA for 48 hours, followed by infection with fluorophore-labeled HPV.FLAG PsVs. To observe fusion events occurring during the early stages of HPV entry after internalization, live-cell imaging was performed at 1.5 hpi, based on our observation of Rabankyrin-5 association with HPV at this time point (Fig. 1G). Representative images in Fig. 2D show examples of HPV particle collision and fusion in siRNA-transfected HeLa cells. In control cells, vesicle ‘a’ collided with vesicles ‘b’ and ‘c’ and underwent multiple fusion events, ultimately forming a large HPV-positive punctum ‘A’ (Fig. 2D, upper panel). In contrast, in Rabankyrin-5 knockdown cells, vesicles ‘a’ and ‘b’ experienced multiple collisions without fusing (Fig. 2D, lower panel). Quantification of fusion events per cell over a 2-minute period revealed that Rabankyrin-5 depletion significantly inhibited fusion (Fig. 2E).

To further determine whether fusion occurs between HPV-carrying early endosomal vesicles or within individual endosomes, we analyzed fusion events in Rab5- and Rabankyrin-5-positive compartments. HeLa cells were co-transfected with Rabankyrin-5-eGFP and mRFP-Rab5. After 48 hours, the cells were infected with HPV16 PsVs labeled with AlexaFluor647. Co-movement of virus particles with mRFP-Rab5 and Rabankyrin-5-eGFP was imaged at 1.5 hpi. Virus-carrying endosomal vesicles positive for both proteins were observed to undergo fusion (Fig. 2F), suggesting successful fusion between vesicles. Co-IP assays using cells co-expressing FLAG-Rab5 and Rabankyrin-5-eGFP revealed increased co-precipitation of both endogenous and exogenous Rabankyrin-5 with L2 proteins in infected cells compared to uninfected controls, indicating that exogenous Rabankyrin-5 and Rab5 form a complex with HPV L2 during infection (Fig. S2F). This molecular interaction further supports the idea that Rabankyrin-5 plays a role in mediating the fusion of HPV-carrying early endosomes.

### Rabanykrin-5 promotes HPV movement along microtubules

How does Rabankyrin-5 facilitate the fusion of HPV-carrying early endosomes? We noted that viral particles exhibited confined movement in Rabankyrin-5-depleted cells, implying that impaired trafficking may contribute to the observed fusion defects and that Rabankyrin-5 is involved in promoting viral transport along microtubules. To test this, HeLa cells were treated with either control siRNA or Rabankyrin-5-targeting siRNA, followed by infection with AlexaFluor488-labeled HPV16 PsVs.

At 1.5 hpi, microtubules were stained to visualize viral trajectories in the microtubule network. Live-cell imaging revealed heterogeneous viral particle dynamics: while some particles were immotile or exhibited slow, drifting behavior, others followed directed paths (Fig. 3). In control siRNA-treated cells, many viral particles displayed robust movement along microtubules, showing directed motions (Fig. 3A, upper panel). In contrast, Rabankyrin-5-depleted cells exhibited a marked increase in confined viral motion (Fig. 3A, lower panel).

**Figure 3.**
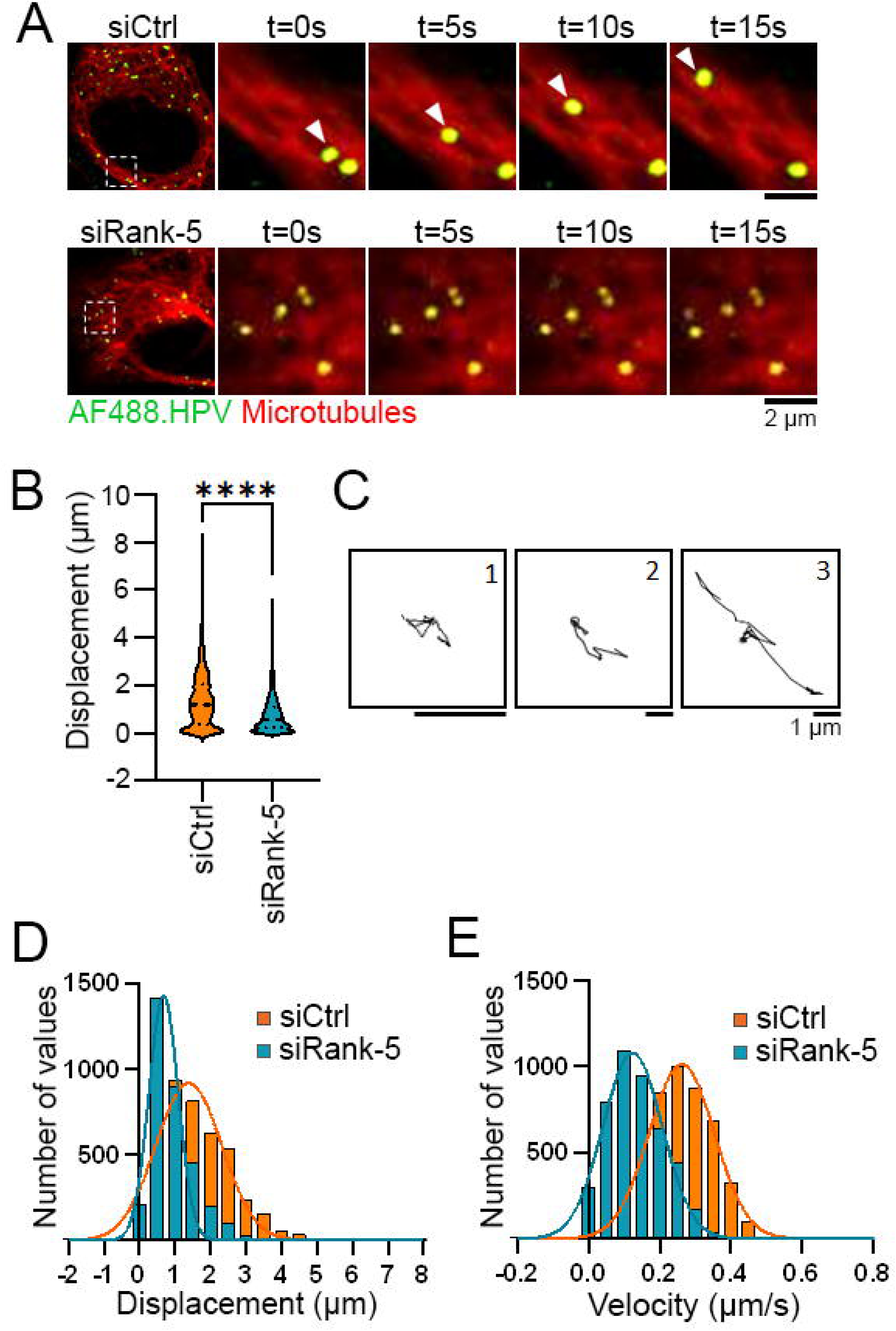
Depletion of Rabankyrin-5 impairs HPV movement along microtubules. (A) HeLa cells were transfected with siRNAs as described in Figure 2A. After 48 hours, cells were infected with AF488.HPV (green) at an MOI of 100. At 1.5 hpi, microtubules were visualized by staining prior to image acquisition. Representative time-lapse insets are shown on the right. Microtubules are pseudocolored in red. Inset scale bar: 2 μm. (B) The displacements of virus particles from their starting to ending positions were analyzed using TrackMate (n = 4500-5000 particles). The graphs show the distribution pattern of the displacements. Statistical significance was assessed using a two-tailed *t*-test with unequal variance, compared to siCtrl-treated cells. ****p < 0.001. (C) Representative trajectories illustrating three distinct modes of virus particle motion with displacements exceeding 0.2 μm are shown: 1) restricted diffusion, 2) random confined motion, and 3) directed movement. Scale bar: 1 μm. (D) Frequency distribution plots of virus displacements greater than 0.2 μm were fitted to a Gaussian distribution. The mean displacement of virus particles was 1.39 ± 0.9 μm in control cells and 0.68 ± 0.4 μm in Rabankyrin-5-depleted cells. (E) Frequency distribution plots of the maximum velocity of each particle during a 2-minute observation period (with displacements > 0.2 μm) were also fitted to a Gaussian distribution. The mean velocity was 0.26 ± 0.09 μm/s in control cells and 0.12 ± 0.08 μm/s in Rabankyrin-5-depleted cells.

To quantitatively analyze the movement patterns of individual HPV particles, we measured their displacement from the starting point to the endpoint of their trajectories. Notably, approximately 30% of particles exhibited minimal movement, with displacements below 0.2 µm (Fig. 3B). Based on this observation, we applied a displacement threshold of 0.2 µm to categorize particle motility: particles with displacements ≤ 0.2 µm were classified as immotile, while those with displacements > 0.2 µm were considered motile. This threshold enabled us to distinguish between stationary particles and those undergoing active transport, allowing for a more focused analysis of the motile population involved in HPV trafficking.

We next extracted viral particle trajectories from time-lapse image series using TrackJ. Among the motile population (displacements > 0.2 µm), three distinct motion types were identified based on established criteria: 1) restricted diffusion, 2) random confined motion, and 3) directed movement (Fig. 3C) (40). In control cells, all three behaviors were observed, including a distinct subset of particles exhibiting highly mobile, directed trajectories consistent with efficient transport along microtubules. In contrast, viral particles in Rabankyrin-5-depleted cells were predominantly restricted to limited diffusion and confined motion, suggesting impaired trafficking.

To quantify this impairment, we compared the displacement of viral particles in control and Rabankyrin-5 knockdown cells. Particles in control cells traveled significantly longer distances than those in knockdown cells. Gaussian fitting of displacement frequency distributions revealed a mean displacement of 1.39 ± 0.9 µm in control cells, compared to 0.68 ± 0.4 µm in Rabankyrin-5-depleted cells (Fig. 3D). We further assessed the effect of Rabankyrin-5 on the velocity of viral transport. Because of the characteristic “move-stop-move” dynamics of HPV particles, we calculated the maximum velocity of each particle during a 2-minute observation period. Gaussian fitting showed a mean maximum velocity of 0.26 ± 0.09 µm/s in control cells, which was reduced to 0.12 ± 0.08 µm/s following Rabankyrin-5 knockdown (Fig. 3E). These findings indicate that Rabankyrin-5 is critical not only for the fusion of virus-containing early endosomes but also for promoting efficient, microtubule-based motility of HPV particles during entry.

### Rabankyrin-5 facilitates HPV transport by connecting dynein to the virus at early endosomes

Based on the observed reduction in the fusion of virus-containing early endosomes and the impaired viral motility in Rabankyrin-5-depleted cells, we propose that HPV transport and early endosomal fusion are coordinated during the early stages of viral entry, with Rabankyrin-5 potentially acting as a specificity factor in dynein-driven transport of HPV along microtubules.

Previous studies have identified a group of proteins that function as dynein adaptors, which interact with both dynein and its cofactor dynactin to mediate the recruitment of dynein to membrane-bound compartments, enabling processive and directed transport along microtubules (19). Since internalized HPV-carrying vesicles enter the endosomal system after endocytosis (7), we first investigated the requirement of known dynein adaptors for infection.

Specifically, we evaluated adaptors previously shown to facilitate dynein association with endosomes carrying cellular cargo, either in a Rab5-dependent or -independent manner (19). HeLa cells were depleted of these adaptors using siRNA, followed by infection with HPV16 PsVs. Strikingly, none of the tested adaptors were essential for infection (Fig. 4A and Fig. S3A). In contrast, the impairment of HPV infection and viral motility observed in Rabankyrin-5-depleted cells suggests a model in which Rabankyrin-5 plays a critical role in recruiting dynein to HPV-containing endosomes.

**Figure 4.**
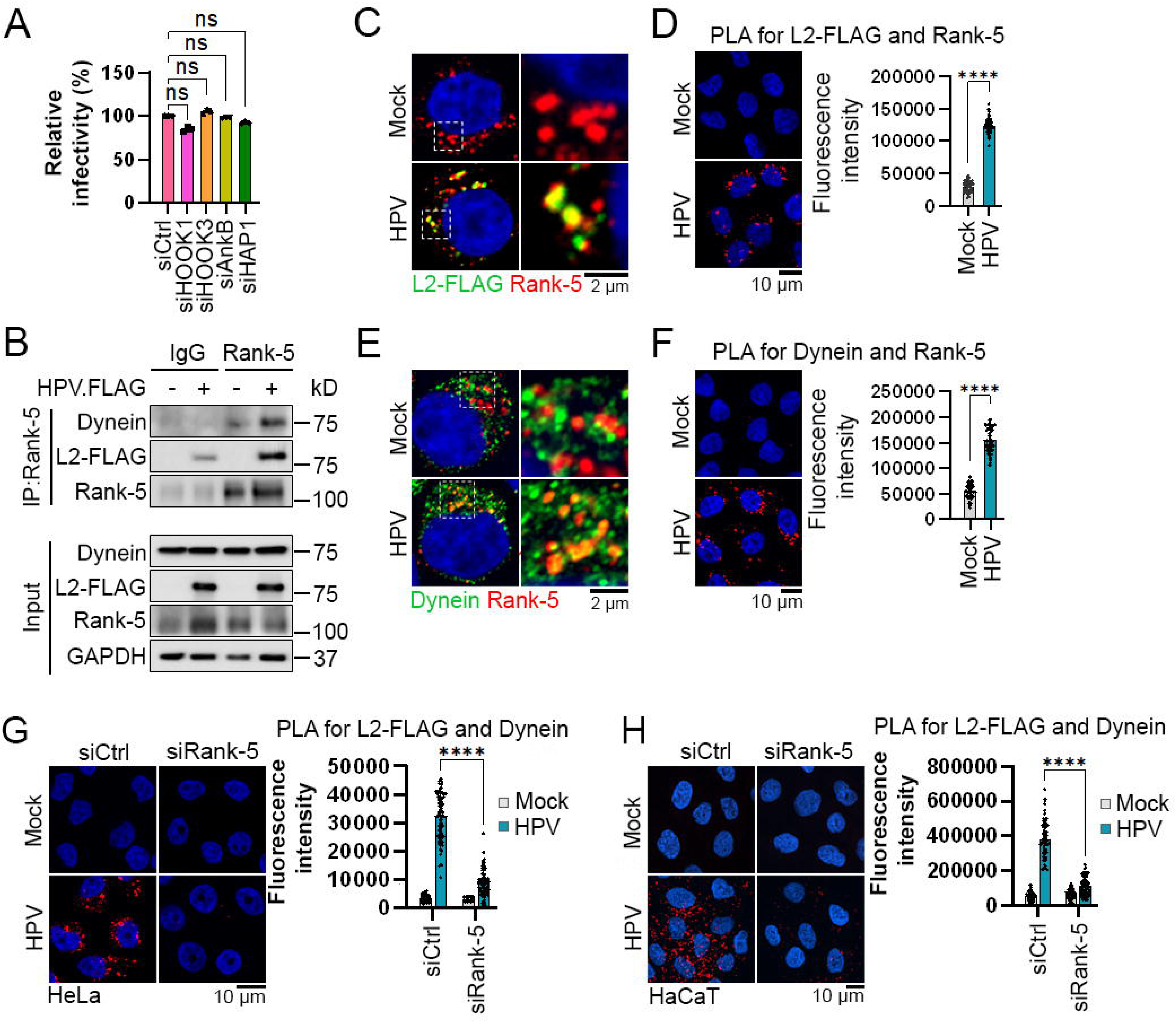
Rabankyrin-5 recruits dynein to HPV by forming a complex with HPV L2 and dynein. (A) HeLa cells were transfected with negative control siRNA (siCtrl) or siRNA targeting known dynein adaptors involved in endosomal cargo transport for 48 hours. The cells were then either mock-infected or infected at an MOI of 2 with HPV.FLAG PsVs containing the HcRed reporter plasmid. At 48 hpi, HcRed fluorescence was measured by flow cytometry to determine relative infectivity, as described in Fig. 1A. Statistical significance was assessed using a two-tailed *t*-test with unequal variance, based on three independent experiments, comparing each condition to siCtrl-treated cells. ns, not significant. (B) HeLa cells were mock-infected (-) or infected (+) with HPV.FLAG for 3 hours. Cell lysates were subjected to immunoprecipitation using an anti-Rabankyrin-5 antibody or Normal Rabbit IgG as a negative control. The precipitated proteins were analyzed by SDS-PAGE and immunoblotting with anti-FLAG, anti-dynein, and anti-Rabankyrin-5 antibodies. (C) HeLa cells were mock-infected or infected with HPV.FLAG PsVs at an MOI of 25. At 6 hpi, cells were stained with anti-FLAG and anti-Rabankyrin-5 antibodies, and fluorescence was visualized by confocal microscopy. Representative images (left) and magnified insets (right) are shown. L2-FLAG (green), Rabankyrin-5 (Rank-5, red), and nuclei (blue) are depicted. Co-localization of L2-FLAG and Rabankyrin-5 is shown in yellow (pseudocolored). Scale bar: 2 μm. (D) HeLa cells were mock-infected or infected with HPV.FLAG PsVs at an MOI of 150. At 6 hpi, a proximity ligation assay (PLA) was performed using antibodies against FLAG and Rabankyrin-5. The PLA signal (red) was visualized, and nuclei were counterstained with DAPI (blue). (Right) Multiple images from three independent experiments were analyzed using Fiji software to quantify PLA fluorescence intensity per cell (n = 80-90 cells). Each dot represents a single cell. Scale bar: 10 μm. Statistical significance was assessed using ANOVA (****p < 0.0001). (E) HeLa cells were mock-infected or infected with HPV.FLAG as in (C), stained at 6 hpi with anti-dynein and anti-Rabankyrin-5 antibodies, and analyzed by confocal microscopy. Dynein (green), Rabankyrin-5 (Rank-5, red), and nuclei (blue) are shown; co-localization appears yellow. Scale bar: 2 μm. (F) Cells treated as in (D) were subjected to PLA using anti-dynein and anti-Rabankyrin-5 antibodies at 6 hpi. PLA signal (red) and nuclei (blue) were visualized, and fluorescence intensity was quantified as in (D). Scale bar: 10 μm. ****p < 0.0001. (G and H) HeLa cells (G) and HaCaT cells (H) were treated with control siRNA or siRNA targeting Rabankyrin-5 for 48 hours, then either left uninfected or infected with HPV.FLAG PsVs at an MOI of 200. At 6 hpi, PLA was performed using antibodies against FLAG and dynein (PLA signal in red), and nuclei were stained with DAPI (blue). (Right) Multiple images from (G) and (H) were analyzed and quantified as described in (D) and (F). ****p < 0.0001. Scale bar: 10 μm.

To test this, we examined Rabankyrin-5 interactions with both HPV and dynein during the early stages of viral entry. We first performed co-IP assays to assess the formation of an L2-Rabankyrin-5-dynein complex. HeLa cells were either mock-infected or infected with HPV.FLAG. At 3 hpi, cell lysates were subjected to immunoprecipitation using an antibody against Rabankyrin-5 or a Normal Rabbit IgG as a negative control. The immunoprecipitates were analyzed by SDS-PAGE and immunoblotted with antibodies against dynein, FLAG, and Rabankyrin-5. Rabankyrin-5 co-precipitated with both dynein and L2-FLAG, indicating that it interacts with both HPV and dynein during viral entry (Fig. 4B).

We next performed immunofluorescence assays to assess the colocalization of the virus with Rabankyrin-5. Mock-infected and HPV.FLAG-infected HeLa and HaCaT cells were stained with antibodies against FLAG and Rabankyrin-5. Consistently, infected cells exhibited clear colocalization of L2-FLAG and Rabankyrin-5 (Fig. 4C and Fig. S3B). We further quantified this association using a PLA assay for Rabankyrin-5 and L2-FLAG, which revealed robust PLA signals in infected cells compared to controls (Fig. 4D and Fig. S3C). To assess the association between Rabankyrin-5 and dynein, we conducted additional immunofluorescence and PLA analyses. These confirmed strong colocalization and significantly increased PLA signals between Rabankyrin-5 and dynein in infected cells (Fig. 4E and F; Fig. S3D and E), further supporting the formation of an L2-Rabankyrin-5-dynein complex during HPV entry. Finally, to determine whether Rabankyrin-5 is essential for this complex formation, we assessed the association between HPV and dynein using a PLA for L2-FLAG and dynein in Rabankyrin-5-depleted HeLa and HaCaT cells. Rabankyrin-5 knockdown resulted in a significant reduction in PLA signals, indicating that the interaction between HPV and dynein is Rabankyrin-5 dependent (Fig. 4G and H).

### Rabanykrin-5 acts as a dynein adaptor during HPV entry by directly binding to HPV L2 protein and dynein

Rab protein effectors play a critical role in enabling dynein to recognize and interact with specific membranous compartments (19). Some of these effectors act as dynein adaptors by directly binding to dynein and facilitating its recruitment to Rab-marked membranes. This interaction is essential for activating dynein motility and supporting efficient microtubule-based transport (19). Although no consensus sequences define dynein adaptors, Rabankyrin-5 contains key structural features common to known adaptors, including a coiled-coil region, a dimerization domain, and a GTPase-binding site that anchors it to membrane compartments (26, 28). Given that Rabankyrin-5 forms a complex with HPV L2 and dynein and promotes HPV transport along the microtubules (Fig. 3 and 4; Fig. S3), we propose that Rabankyrin-5 functions as a dynein adaptor. Specifically, it may serve as a molecular bridge, simultaneously interacting with HPV and dynein to promote the recruitment and activation of dynein for retrograde transport of the virus.

To assess whether Rabankyrin-5 can directly bind to both the virus and dynein as a dynein adaptor, we first examined its direct interaction with the C-terminal region of the L2 protein. Since a segment of the L2 C-terminus becomes exposed to the cytosol following membrane protrusion (13), we performed a pull-down assay using purified GST or GST-tagged Rabankyrin-5 in combination with purified GFP or a GFP-L2-C fusion protein containing the final 40 residues of HPV16 L2. After incubation, GST or GST-Rabankyrin-5 was pulled down using glutathione beads, and bound proteins were analyzed by SDS-PAGE followed by immunoblotting. As shown in Fig. 5A, GST-Rabankyrin-5, but not GST alone, pulled down the GFP-L2-C fusion protein, indicating a direct interaction between Rabankyrin-5 and the L2 C-terminus. To assess dynein binding, a similar pull-down assay was performed using purified dynein complex. Dynein precipitated with GST-Rabankyrin-5 but not with GST alone, confirming a direct interaction between Rabankyrin-5 and dynein (Fig. 5B). Together with our previous findings, these results demonstrate that Rabankyrin-5 directly binds both the HPV L2 protein and the dynein complex.

**Figure 5.**
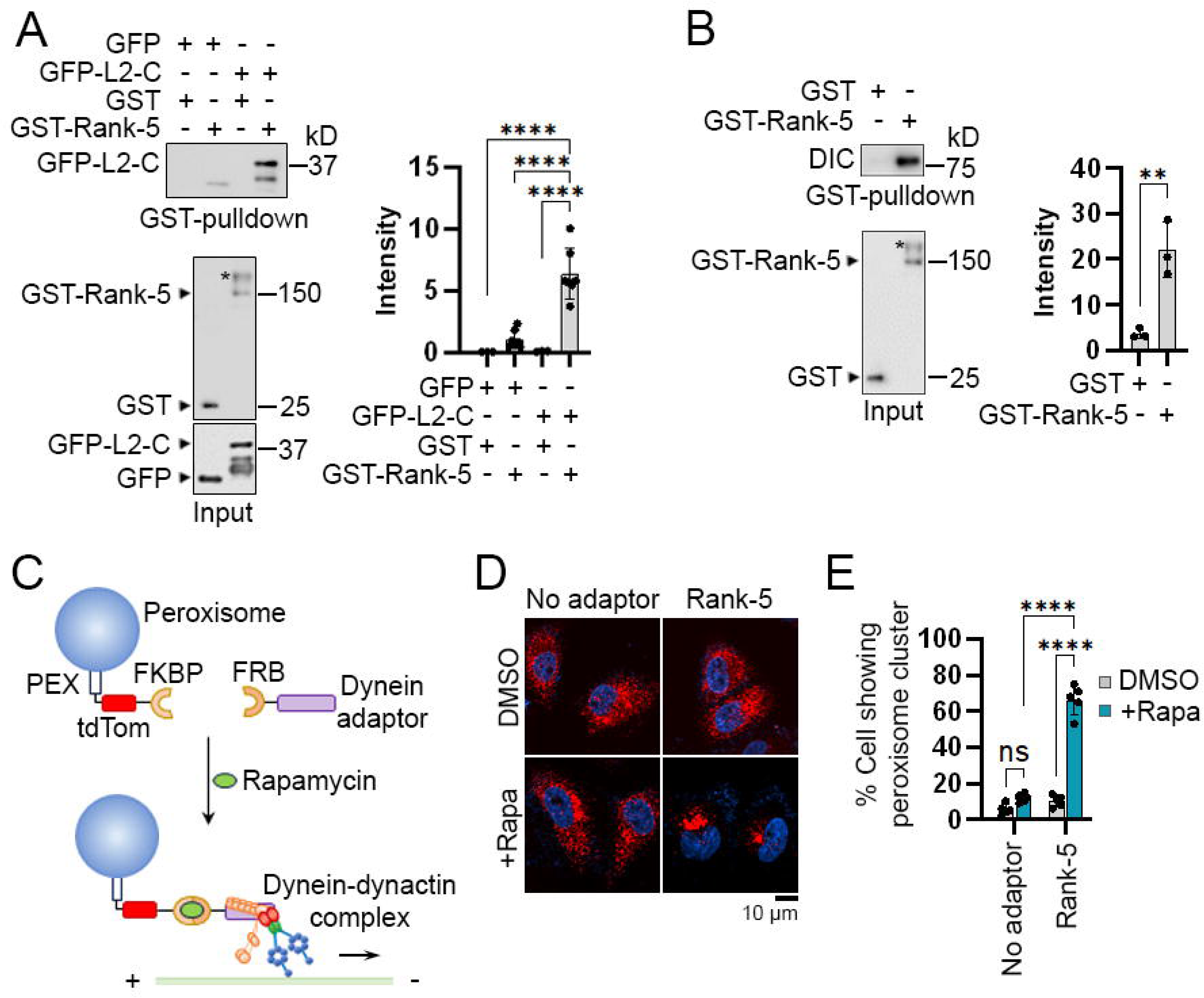
Rabankyrin-5 interacts with HPV L2 and dynein and serves as a dynein adaptor protein. (A) Purified GST or GST-tagged Rabankyrin-5 (GST-Rank-5) bound to glutathione beads was incubated with either purified His-tagged GFP or His-tagged GFP-L2-C fusion protein, which contains the C-terminal 40 amino acids of L2. After the pulldown, bound proteins were analyzed by SDS-PAGE and detected with an anti-GFP antibody (top panel). The bottom panel shows input GST and GST-Rank-5 detected with an anti-GST antibody, and GFP and GFP-L2-C detected with an anti-His antibody. The band around 300 kDa is likely Rabankyrin-5 dimers, indicated by asterisks (*). (Right) Band intensities from at least three independent experiments were quantified and plotted. Data are presented as mean ± SD. ****p < 0.0001. (B) GST or GST-Rank-5, as in (A), was incubated with or without purified cytoplasmic dynein-1 complex. After the pulldown, bound proteins were analyzed by SDS-PAGE and immunoblotted with an anti-dynein antibody (top panel). The bottom panel shows input GST and GST-Rank-5 proteins detected with anti-GST antibodies. (Right) Band intensities from three independent experiments were quantified. Data are presented as mean ± SD. **p < 0.01. (C) Schematic of the inducible peroxisome trafficking assay. Cells are transfected to express tdTomato-FKBP fused to a peroxisome-targeting sequence (PEX), together with a construct encoding a candidate dynein adaptor protein fused to an FRB domain. Rapamycin treatment induces FKBP-FRB dimerization, recruiting the dynein-dynactin complex to peroxisomes via the dynein adaptor and initiating retrograde transport along microtubules. (D) HeLa cells were co-transfected with a construct expressing PEX3-tdTomato-FKBP and a construct expressing either FRB alone (No adaptor) or Rabankyrin-5-FRB (Rank-5). After 24 hours, cells were treated with either DMSO or rapamycin (+Rapa), then fixed and imaged. Scale bar: 10 μm. (E) Quantification of the percentage of cells showing strong peroxisome clustering near the nucleus. Data represent mean ± SD from three independent experiments. ****p < 0.0001; ns, not significant.

Because most dynein adaptors are known to function as activators of dynein-driven transport (19), we tested whether Rabankyrin-5 can act as a dynein activator using a cell-based, inducible peroxisome trafficking assay. In this system, cells co-express two fusion constructs: one comprising the peroxisomal targeting sequence PEX, the fluorescent protein tdTomato, and the FK506-binding protein domain (PEX-tdTM-FKBP), and another containing the candidate adaptor protein fused to the FKBP12-rapamycin-binding (FRB) domain (41–43). Upon treatment with a rapamycin analog, the FKBP and FRB domains dimerize, thereby recruiting the candidate protein to peroxisomes. If the candidate functions as an activating dynein adaptor, it will engage the dynein-dynactin complex and promote their minus-end-directed transport along microtubules. Enhanced peroxisome relocation to the perinuclear region under these conditions thus provides a quantitative readout of dynein activation by the candidate protein (Fig. 5C).

We constructed expression plasmids encoding either FRB alone or Rabankyrin-5-FRB, which were co-expressed with the fusion protein PEX-tdTM-FKBP. In control cells expressing FRB alone, with or without rapamycin, tdTomato-marked peroxisomes were distributed throughout the cytoplasm. A similar distribution was observed in cells transfected with Rabankyrin-5-FRB in the absence of rapamycin. However, upon rapamycin treatment, cells expressing Rabankyrin-5-FRB exhibited peroxisome clustering near the nuclear region in approximately 70% of cells, suggesting that Rabankyrin-5-FRB possesses intrinsic dynein-activating activity (Fig. 5D and E).

Together with the observed defects in viral motility (Fig. 3), these findings suggest that the impaired movement of HPV particles along microtubules in Rabankyrin-5-depleted cells results from disrupted interactions between the virus and dynein. Rabankyrin-5 may function to recruit dynein complexes to HPV-containing early endosomes, facilitating their directional transport along microtubules. This recruitment likely promotes proper spatial positioning of the vesicles and primes them for fusion by guiding their approach toward target membranes. Disruption of the association between the virus and dynein in Rabankyrin-5-depleted cells may lead to mispositioned virus-carrying endosomes and, consequently, a reduction in fusion events (Fig. 6). These findings reveal a previously unrecognized role for Rabankyrin-5 in coordinating HPV trafficking and endosomal fusion during early stages of infection.

**Figure 6.**
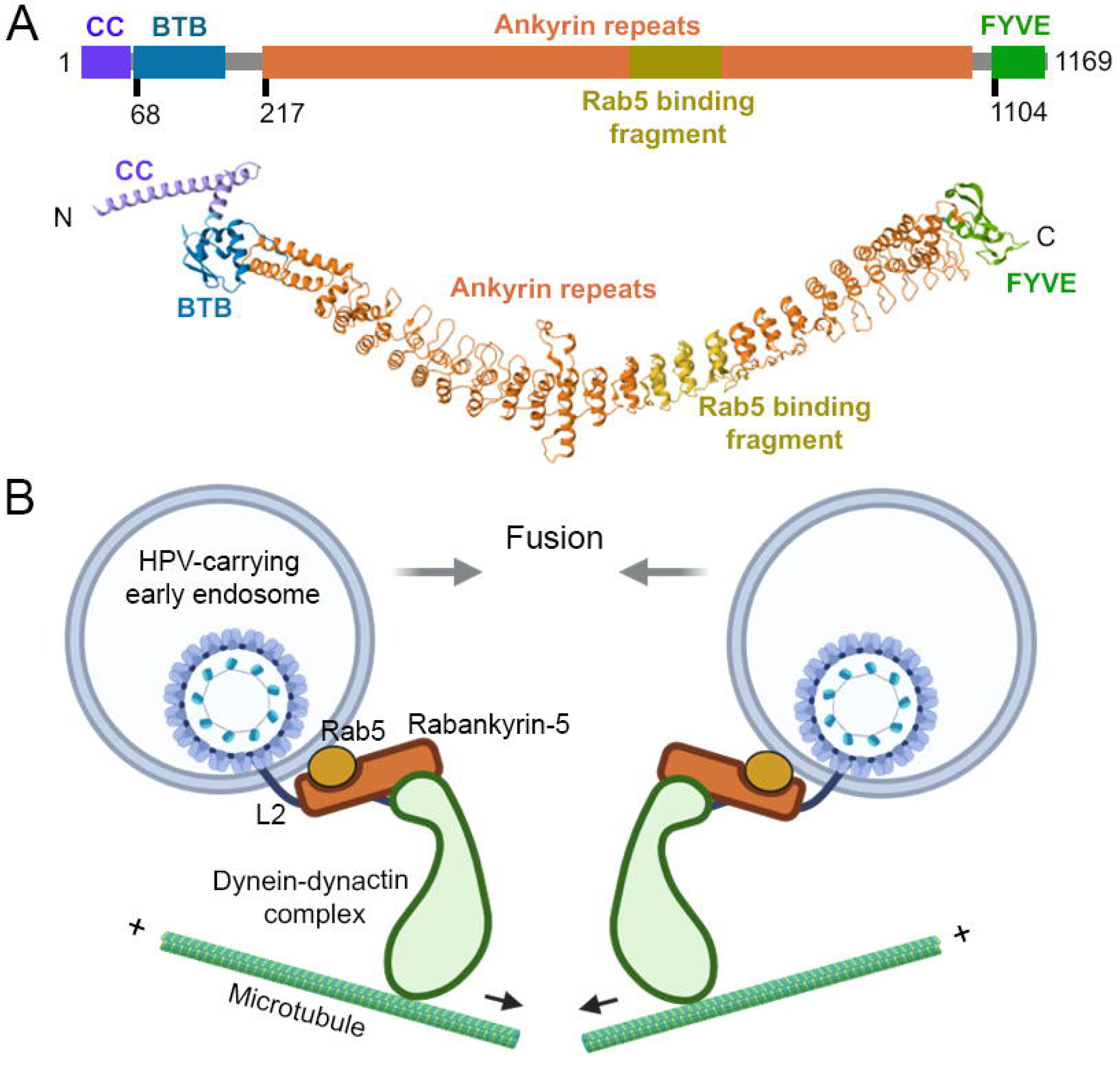
Proposed model of Rabankyrin-5-mediated fusion of HPV-carrying early endosomes. (A) Schematic diagram of the domain organization in human Rabankyrin-5 within the context of the full-length protein (top). Rabankyrin-5 consists of four distinct domains: a coiled-coil (CC) domain, a Broad-Complex, Tramtrack, and Bric-à-brac (BTB) domain, middle ankyrin repeats containing a Rab5 binding fragment, and a PI(3)P-binding FYVE domain. Domain sizes are not to scale. Structured domains predicted by AlphaFold2 are shown (bottom). (B) Proposed model of Rabankyrin-5-mediated fusion of HPV-carrying early endosomes and their microtubule-based transport following endocytosis. Upon cytosolic exposure of L2 from the early endosomal membrane, the HPV L2 protein binds to the Rab5-associated Rabankyrin-5 complex, which is linked to the dynein motor machinery. This complex forms a specific interaction between the virus-containing early endosome and dynein-dynactin complex, enabling directed transport along microtubules, facilitating proper positioning of the early endosomes, and promoting fusion of these endosomes during viral entry. Figure created with BioRender.com.

## Discussion

Following endocytosis, newly internalized, cargo-enriched endocytic vesicles fuse to form early endosomes, which subsequently mature into late endosomes (1, 2). The successful fusion of these vesicles depends on the precise coordination of Rab GTPases, their effectors, and SNARE proteins (22, 29, 44). While SNAREs act as molecular machinery that drive membrane priming, docking, and the physical fusion of lipid bilayers, Rab GTPases and their effectors define compartment identity and serve as targeting and tethering factors that mediate vesicle capture and positioning for fusion (21, 22, 39). Rabankyrin-5, a Rab5 effector, was found to be essential for HPV infection (Fig. 1B) (25). Although its role in membrane fusion is not well characterized (26, 29), depletion of Rabankyrin-5 impairs the fusion of HPV-carrying early endosomes. Notably, its depletion also disrupts viral motility (Fig. 2 and 3), suggesting that the early endosomal fusion may be coordinated with the trafficking of these endosomes during HPV entry. Given that the motor protein dynein interacts with cargo and drives their transport along microtubules, we hypothesized that dynein is involved in the fusion in concert with Rabankyrin-5. Our co-IPs, pull-down assays, and imaging studies demonstrate that Rabankyrin-5 directly binds both the HPV L2 capsid protein and the dynein complex (Fig. 4, 5A, and 5B). Furthermore, in a cell-based inducible organelle relocation assay, Rabankyrin-5 appears to activate the dynein transport complex (Fig. 5C, D, and E). These findings suggest that Rabankyrin-5 acts as a novel dynein adaptor, linking dynein to virus-containing early endosomes and facilitating microtubule-based transport that promotes endosomal fusion.

Another Rab5 effector, EEA1, has been shown to promote early endosome fusion (21, 22, 29, 45). Recent studies suggest that EEA1, a long coiled-coil protein, undergoes large-scale conformational changes upon binding to Rab5 and phosphatidylinositol 3-phosphate (PI(3)P), a lipid enriched in early endosomal membranes. Through these interactions, EEA1 acts as a long-range tether that pulls vesicles toward the target membrane to initiate docking and fusion (21, 22, 45). Interestingly, depletion of EEA1 does not significantly affect HPV PsV infection in HeLa and HaCaT cells, consistent with our observation that EEA1 does not co-precipitate with HPV L2 proteins (data not shown). Given that EEA1 is unevenly distributed across early endosomal membranes, it is possible that EEA1 is present on HPV-carrying early endosomes without contributing to their fusion, or that a subset of early endosomes is EEA1-positive but does not participate in the fusion of HPV-containing compartments. In contrast, Rabankyrin-5, which also possesses Rab5 and PI(3)P binding domains (26), interacts with the HPV L2 protein and mediates the fusion of virus-containing early endosomes (Fig. 2 and 6A). These findings suggest that HPV may exploit the host fusion machinery in a manner distinct from cellular cargo, preferentially utilizing Rabankyrin-5, despite its weaker tethering capability compared to EEA1 for cellular cargo (21, 29, 33), to avoid competition for fusion sites and increase its chances of successful infection.

The endosomal system is highly dynamic, relying on the coordinated interactions of specific proteins and lipids to ensure precise membrane targeting and tethering for vesicle fusion. During HPV entry, the L2 protein likely contributes to this specificity by facilitating the assembly of a membrane-targeting complex with Rabankyrin-5 at the early endosomal membrane (Fig. 4 and 5). This complex not only defines the fusion site but also mediates the recruitment of dynein (Fig. 4). Dynein recruitment, in turn, facilitates the microtubule-based transport of virus-containing endosomes, promoting their proper positioning and directing them toward target membranes for subsequent fusion.

One possible model is that the L2 protein first penetrates the endosomal membrane and then engages Rabankyrin-5, thereby initiating dynein recruitment to the vesicle. Alternatively, dynein may already be associated with, or in close proximity to, Rabankyrin-5-positive endosomes, and the exposure of L2 may stabilize or enhance the formation of a tethering complex by interacting with Rabankyrin-5 and recruiting additional dynein molecules. This model is supported by increased co-precipitation and colocalization of dynein with Rabankyrin-5 following HPV infection, suggesting that infection enriches dynein at sites where L2 and Rabankyrin-5 are present (Fig. 4). In either scenario, the interaction between L2 and Rabankyrin-5 likely ensures the specificity of dynein attachment to virus-containing early endosomes, coordinating trafficking with fusion.

Dynein is responsible for transporting a wide range of membrane-bound cargoes. Its adaptor proteins interact with dynein and mediate cargo selectivity across different cellular compartments and organelles (19). Our study shows that Rabankyrin-5 functions as a dynein adaptor, coupling the trafficking and fusion of virus-containing early endosomes following endocytosis. Recent findings suggest that Ran-binding protein 10 (RanBP10) and karyopherin alpha 2 (KPNA2) form a complex that may serve as a dynein adaptor, facilitating the nuclear delivery of the HPV genome during mitosis (35). Additionally, the known dynein adaptor BICD2 mediates HPV transport from endosomes through the Golgi (36, 46). These findings indicate that distinct dynein adaptors are engaged at different stages of HPV entry.

The conversions of Rab proteins, their effectors, and phosphoinositides regulate specific protein-protein and protein-lipid interactions (1, 19, 21, 29). This mechanism may facilitate the timely release of dynein from the virus at its intended intracellular destination, allowing subsequent association with new adaptor proteins to support continued trafficking toward downstream compartments. Rabankyrin-5 contains a FYVE domain that binds PI(3)P (26, 28) (data not shown). This interaction may play a role in regulating dynein dissociation, especially during the conversion of PI(3)P to PI(3,5)P₂ at the late endosome, a transition that also involves the replacement of Rab5 with Rab7 (1). Moreover, stage-specific interactions between the L2 protein and various Rab GTPases and their effectors during infection may further contribute to the regulation of dynein recruitment and cargo specificity throughout HPV intracellular trafficking.

Collectively, our data indicate that Rabankyrin-5 plays a key role in the fusion of virus-carrying early endosomes. It recruits the dynein complex to these endosomes by interacting with both the C-terminal segment of the HPV L2 capsid protein and dynein. This recruitment facilitates microtubule-based transport of the virus-carrying endosomes, enhancing their encounter and subsequent fusion (Fig. 6). Thus, Rabankyrin-5 couples dynein-mediated transport with endosomal fusion to enable efficient HPV entry. Our findings suggest that endosomal trafficking and fusion are functionally linked, highlighting the importance of intracellular transport along the microtubule network in regulating fusion during the formation of early endosomes.

## Materials and Methods

### Cells, Plasmids, and Viruses

HeLa (ATCC, catalog no. CCL-2), HeLa S3 (ATCC, catalog no. CCL-2.2), and 293TT cells (ATCC, catalog no. CRL-3467) were obtained from the American Type Culture Collection (ATCC). HaCaT cells (catalog no. T0020001) were purchased from AddexBio Technologies. Human foreskin keratinocytes (HFK) cells were a gift from Dr. Xuefeng Liu (The Ohio State University). HeLa, HaCaT, and 293TT cells were cultured at 37°C in DMEM (Corning) with HEPES and L-glutamine, supplemented with 10% fetal bovine serum (FBS) and 100 units/mL penicillin streptomycin, in 5% CO_2_. HFK cells were cultured at 37°C in complete Keratinocyte-SFM medium supplemented with rEGF (0.2 ng/mL), BPE (20 μg/mL), and gentamicin (5 μg/mL).

The plasmid p16sheLL, which expresses both L1 and L2 for HPV16 pseudovirus production, was a gift from John Schiller (Addgene plasmid #37320). The pCAG-HcRed reporter plasmid was purchased from Addgene (plasmid #11152). HPV16 PsVs containing a 3 × FLAG tag or HA tag at the C-terminus of L2 (p16sheLL.FLAG and p16sheLL.HA, respectively) were gifts from Dr. Daniel DiMaio (Yale University) (47). The Rabankyrin-5 coding sequence was purchased from GenScript and cloned into pcDNA3.1(+)eGFP to generate pcDNA3.1(+)Rank-5-eGFP, and into pET-41a(+) to generate pET-41a(+)GST-Rank-5, which contains GST at the N-terminus of Rabankyrin-5. The plasmids mRFP-Rab5 and FLAG-Rab5A were gifts from Drs. Ari Helenius (Addgene plasmid #14437) and Qing Zhong (Addgene plasmid #28043), respectively. For the peroxisome relocation assay, pBa.PEX3-tdTomato-FKBP and HA-BICD2(1–500)-FRB were gifts from Drs. Marvin Bentley (Addgene plasmid #193941) and Lukas Kapitein (Addgene plasmid #174648). HA-Rank-5-FRB was generated by replacing BICD2 in HA-BICD2(1–500)-FRB with Rabankyrin-5. To construct HA-FRB, BICD2 was removed from HA-BICD2(1–500)-FRB, and the resulting plasmid was used as a negative control in the assay. pET-41a(+)-GFP and pET-41a(+)-GFP-L2-C were constructed by synthesizing the DNA sequences encoding GFP or GFP-L2-C and replacing the GST coding region in pET-41a(+).

HPV16 PsVs with a 3 × FLAG tag or HA tag at the C-terminus of L2 (HPV.FLAG and HPV.HA, respectively) were assembled by co-transfecting p16sheLL.FLAG or p16sheLL.HA with pCAG-HcRed (as the reporter plasmid) into 293TT cells. Packaged PsVs were purified using density gradient centrifugation in OptiPrep, as described (38). Briefly, following transfection, cells were collected and lysed in buffer containing 0.5% Triton X-100, 10 mM MgCl₂, 5 mM CaCl₂, and 0.5 U/mL RNase A. The lysate was then subjected to density gradient centrifugation in OptiPrep at 50,000 rpm in an SW55 Ti rotor for 4 hours. PsV quality was assessed by Coomassie Brilliant Blue staining for HPV16 L1 and L2 after SDS-PAGE. The virus titer was determined by flow cytometry, based on fluorescent reporter protein expression in HeLa S3 cells 48 hours post-infection.

For live cell imaging, purified HPV.FLAG PsVs were labeled following the manufacturer’s protocol of the Protein Labeling Kit (Invitrogen) (Table 1). Briefly, 1 mg of virus in 0.5 mL PBS was incubated with a vial of reactive dye for 1 hour at room temperature. Purification Spin Columns were used to remove non-conjugated fluorescent dyes. The degree of labeling (DOL) was determined by spectrophotometry.

**Table 1.** Antibodies, siRNA, reagents, and commercial assays.

| **Antibodies** |  |  |
| --- | --- | --- |
| Mouse anti-L1 Protein of Human Papilloma Virus clone CAMVIR-1 (RUO) | BD Biosciences | Cat#554171 |
| Rabbit monoclonal antibody EEA1 (C45B10) | Cell Signaling Technology | Cat#3288 |
| DYKDDDDK Tag (D6W5B) Rabbit mAb | Cell Signaling Technology | Cat#14793 |
| DYKDDDDK Tag (9A3) Mouse mAb | Cell Signaling Technology | Cat#8146 |
| Rabbit monoclonal antibody EEA1 (C45B10) | Cell Signaling Technology | Cat#3288 |
| Mouse monoclonal anti-GAPDH (6C5) | Santa Cruz Biotechnology | Cat#sc-32233 |
| Mouse monoclonal anti-GFP (B-2) | Santa Cruz Biotechnology | Cat#sc-9996 |
| Donkey anti-Mouse IgG (H+L) Highly Cross-Adsorbed Secondary Antibody, Alexa Fluor 488 | Thermo Fisher | Cat#A21206 |
| Donkey anti-Rabbit IgG (H+L) Highly Cross-Adsorbed Secondary Antibody, Alexa Fluor 555 | Thermo Fisher | Cat#A31572 |
| Donkey anti-Rabbit IgG (H+L) Highly Cross-Adsorbed Secondary Antibody, Alexa Fluor 568 | Thermo Fisher | Cat#A10042 |
| Donkey anti-Rabbit IgG (H+L) Highly Cross-Adsorbed Secondary Antibody, Alexa Fluor 647 | Thermo Fisher | Cat#A32795 |
| Dynein Monoclonal Antibody (74.1) | Thermo Fisher | Cat#MA1-070 |
| ANKFY1 Polyclonal Antibody | Thermo Fisher | Cat#PA5-66801 |
| RAB5A Monoclonal Antibody (2E8B11) | Thermo Fisher | Cat#14-9711-82 |
| RAB5 Polyclonal Antibody | Thermo Fisher | Cat#PA5-29022 |
| GST Tag Monoclonal Antibody (8-326) | Thermo Fisher | Cat#MA4-004 |
| 6x-His Tag Monoclonal Antibody (HIS.H8) | Thermo Fisher | Cat#MA1-21315 |
| Monoclonal ANTI-FLAG® M2 antibody | Sigma | Cat#3165 |
| Normal Rabbit IgG | Sigma | Cat#NI01 |
| Monoclonal ANTI-FLAG® M2-Peroxidase (HRP) antibody | Sigma | Cat#8592 |
| **siRNA** |  |  |
| siGENOME Non-Targeting siRNA Pool | Dharmacon | Cat# D-001206-13-05 |
| siGENOME Human ANKFY1 (51479) | Dharmacon | Cat# D-013161-02-0005 |
| siGENOME Human RAB5A (5868) | Dharmacon | Cat# MQ-004009-00-0005 |
| siGENOME Human HOOK1 (51361) | Dharmacon | Cat# M-016845-01-0005 |
| siGENOME Human HOOK3 (84376) | Dharmacon | Cat# M-013558-00-0005 |
| siGENOME Human HAP1 (51361) | Dharmacon | Cat# M-011560-01-0005 |
| siGENOME Human ANK2 (287) | Dharmacon | Cat# M-008417-02-0005 |
| **Chemicals and Proteins** |  |  |
| Rapamycin | Thermo Fisher | AAJ62473MF |
| Hoechst33342, Trihydrochloride, Trihydrate | Thermo Fisher | Cat#H3570 |
| Halt Protease and Phosphatase Inhibitor Single-Use Cocktail, EDTA-Free | Thermo Fisher | Cat#78443 |
| Cytoplasmic Dynein Motor Protein | Cytoskeleton, Inc | Cat#CS-DN01 |
| **Commercial Assays** |  |  |
| Duolink® In Situ PLA Probe Anti-Mouse PLUS | Sigma | Cat#DUO92001 |
| Duolink® In Situ PLA Probe Anti-Rabbit MINUS | Sigma | Cat#DUO92005 |
| Duolink® In Situ Detection Reagents Red | Sigma | Cat#DUO92008 |
| Anti-FLAG M2 affinity gel | Sigma | Cat#A2220 |
| iScript™ cDNA Synthesis Kit | Bio-rad | Ca#1708891 |
| iQ™SYBR Green Supermix | Bio-rad | Cat#1708882 |
| SiR-Tubulin Kit | Cytoskeleton, Inc | Cat#CY-SC002 |
| Ni-NTA agarose | Qiagen | Cat#30210 |
| Lipofectamine™ 2000 Transfection Reagent | Thermo Fisher | Cat#11668027 |
| Lipofectamine™ RNAiMAX Transfection Reagent | Thermo Fisher | Cat#13778100 |
| Pierce® Glutathione Agarose | Thermo Fisher | Cat#16100 |
| BCA Protein Assay Kit | Thermo Fisher | Cat#23225 |
| Fluorescent Protein Labeling Kits, Alexa Fluor 488 | Thermo Fisher | Cat#A10235 |
| Fluorescent Protein Labeling Kits, Alexa Fluor 647 | Thermo Fisher | Cat#A20173 |

### Infectivity Assay

To assess the infectivity of HPV PsVs in knockdown cells, 2.5 × 10^4^ HeLa S3, 1.5 × 10^4^ HaCaT, or 3 × 10^4^ HFK cells were seeded in 24-well plates and transfected with 10 nM negative control siRNA or siRNA targeting Rab5A, Rabankyrin-5, or other adaptor proteins using RNAiMAX (Invitrogen) (Table 1). After 48 hours, cells were infected with HPV.FLAG at an MOI of 2. At 48 hours post-infection (hpi), cells were analyzed by flow cytometry to determine the mean fluorescence intensity (MFI). Infectivity in control siRNA-treated cells was set to 100%, and infectivity in targeted siRNA-treated cells was normalized to the control.

### Reverse transcription qPCR

5 × 10^4^ HeLa S3, 3 × 10^4^ HaCaT, or 6 × 10^4^ HFK cells were seeded in 12-well plates and transfected with 10 nM negative control siRNA or siRNA targeting Rab5A, Rabankyrin-5, or other adaptor proteins using RNAiMAX (Invitrogen). After 48 hours, cells were harvested, and total RNA was isolated using the RNeasy Mini Kit (Qiagen) according to the manufacturer’s instructions. One microgram of RNA from each condition was reverse-transcribed into cDNA using the iScript cDNA Synthesis Kit (Bio-Rad) according to the manufacturer’s instructions. qPCR was performed using iQ SYBR Green Supermix (Bio-Rad) in the CFX Real-Time PCR Detection System (Bio-Rad). Primers used in this study are listed in Table 2.

**Table 2.**
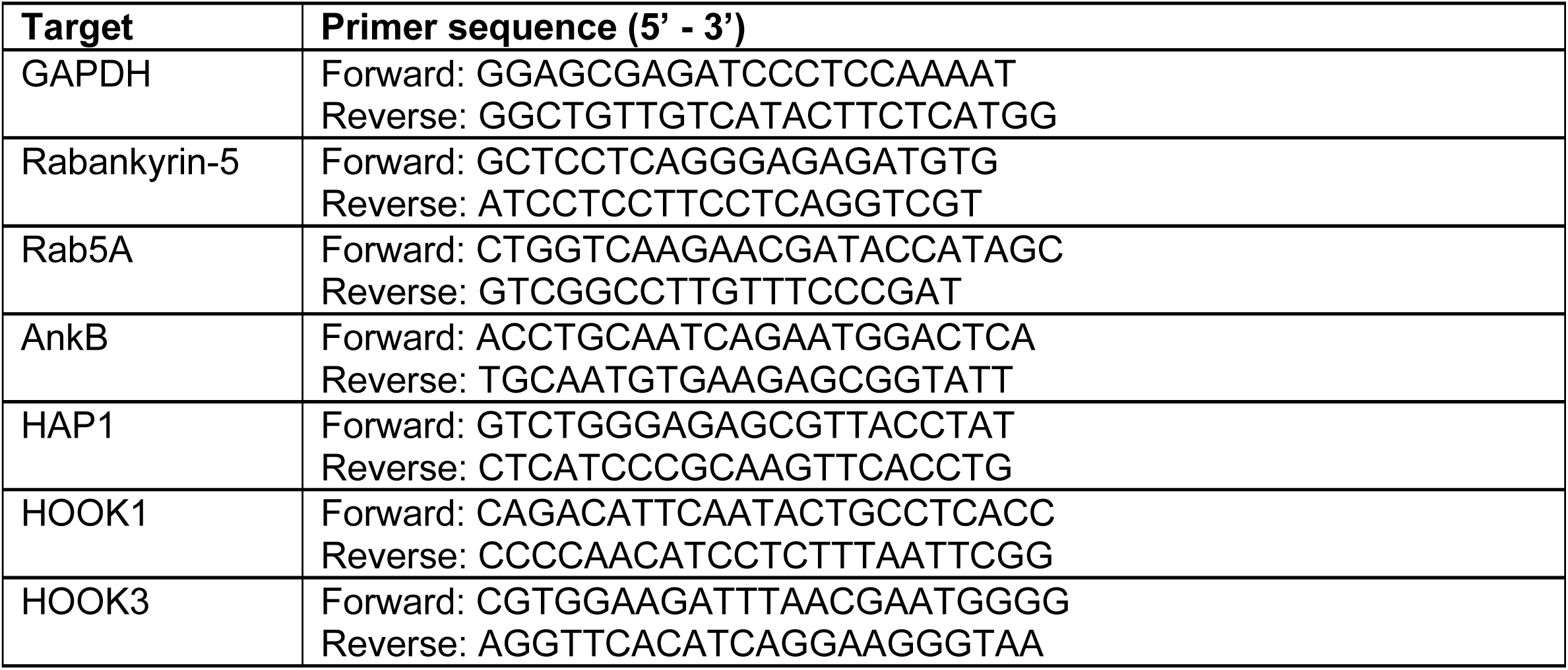
DNA primers used for qPCR analyses.

### Immunofluorescence microscopy and proximity ligation assay

2.5 × 10^4^ HeLa S3 or 1.5 × 10^4^ HaCaT cells were seeded on glass coverslips in 24-well plates and either mock-infected or infected with HPV.FLAG at an MOI of 25. At designed hpi, cells were fixed with 4% paraformaldehyde, permeabilized with 1% saponin, and incubated with primary antibodies against FLAG, EEA1, TGN46, Rab5, Rabankyrin-5, or dynein. Cells were then incubated with AlexaFluor-conjugated secondary antibodies. The slides were mounted, and images were acquired using a Nikon Ti2 spinning disk confocal microscope. All antibodies used in this study are listed in Table 1.

For the proximity ligation assay (PLA), HeLa S3 or HaCaT cells were infected with HPV.FLAG at an MOI of 150. Infected cells were fixed at 6 hpi with 4% paraformaldehyde, permeabilized with 1% saponin, and incubated with paired primary antibodies (one from mouse and one from rabbit). PLA was performed using Duolink reagents according to the manufacturer’s instructions (Sigma). Briefly, following primary antibody staining, cells were incubated in a humidified chamber with PLA probes diluted 1:5, followed by ligation and amplification with a fluorescent substrate at 37°C. Nuclei were stained with DAPI using BD fluorescence mounting medium, and images were acquired using a Nikon Ti2 spinning disk confocal microscope. For quantification, approximately 200 nuclei per sample were imaged.

Images were processed and quantitatively analyzed with Fiji to measure the total fluorescence intensity per cell. All imaging experiments were performed independently in triplicate. To assess Rabankyrin-5 dependence, 1 × 10^4^ HeLa S3 or HaCaT cells were seeded on coverslips in 24-well plates and transfected with 10 nM siRNA for 48 hours before infection with HPV.FLAG at an MOI of 200. Infected cells were fixed and permeabilized as described above at 6 hpi, followed by PLA assessment. Cells transfected with non-target siRNA served as the negative control.

### Co-immunoprecipitation

1.5 × 10^6^ HeLa cells were seeded into 6 cm^2^ dishes. After 16 hours, cells were either mock-infected or infected with HPV.FLAG at an MOI of 25. At designated time points, cells were washed twice with PBS and cross-linked with 1.5 mM DSP [dithiobis(succinimidyl propionate)] for 30 minutes at room temperature. The reaction was quenched using 100 mM Tris-HCl (pH 7.4) for 15 minutes at room temperature. Cells were then washed with cold PBS and lysed in 400 μL of lysis buffer (20 mM HEPES [pH 7.4], 50 mM NaCl, 5 mM MgCl₂, 1 mM DTT) with 0.5% Triton X-100 supplemented with 1 × Halt protease and phosphatase inhibitor cocktail. Lysates were centrifuged at 16,100 g for 10 minutes at 4 °C. Supernatants were incubated with 40 μL anti-FLAG affinity gel and gently rocked overnight at 4 °C. The beads were then washed with lysis buffer containing 0.1% Triton X-100, and bound proteins were eluted in 40 μL of 4 × SDS sample buffer by heating at 95 °C for 10 minutes. Samples were analyzed by SDS-PAGE and immunoblotting using specific antibodies. For Rabankyrin-5 IP, mock infected or infected cells were lysed in RIPA buffer supplemented with 1 × Halt protease and phosphatase inhibitor cocktail, followed by centrifugation at 16,100 g for 10 minutes at 4 °C. Supernatants were incubated with either a Rabankyrin-5 antibody or an equal amount of Normal Rabbit IgG control and gently rocked for 3 hours at 4 °C. Protein A/G agarose beads (Santa Cruz) were then added and incubated for 60 minutes at 4 °C, followed by four washes with RIPA buffer. Bound proteins were eluted by boiling in 4 × SDS sample buffer containing 2-mercaptoethanol at 95 °C for 10 minutes and analyzed by immunoblotting. In Fig. S2F, 2.5 × 10^5^ HeLa cells were plated in 6 cm² dishes. After 16 hours, cells were either mock-transfected or transfected with plasmids encoding FLAG-Rab5A and Rabankyrin-5-eGFP for 48 hours, then mock-infected or infected with HPV.HA at an MOI of 25. Cell lysates were subjected to FLAG IP as described above.

### Protein purification

The plasmid expressing GFP-L2-C or GST-Rabankyrin-5 was transformed into Rosetta (DE3) competent cells (Sigma). Bacterial cultures were grown to an OD_600_ of ∼0.4 and induced with 0.2 mM isopropyl β-D-1-thiogalactopyranoside (IPTG) at 25 °C for 18 hours. To purify GST-Rabankyrin-5, bacteria were harvested and lysed in Lysis buffer (20 mM HEPES, 150 mM NaCl, 20 mM EDTA, 5 mM DTT, 1.5% Triton X-100, 1% Sarkosyl) supplemented with 20 units/mL DNase I, 0.5 mg/mL lysozyme, and 1 mM PMSF. Lysates were incubated with a pre-equilibrated slurry of glutathione beads in Wash buffer (20 mM HEPES, 150 mM NaCl, 20 mM EDTA, 5 mM DTT, 1% Triton X-100) for purification. Bound proteins were eluted using Elution buffer (20 mM HEPES, 150 mM NaCl, 20 mM EDTA, 5 mM DTT, 0.1% Triton X-100, 40 mM reduced glutathione, 1% Sarkosyl), then dialyzed into buffer containing 50 mM Tris-HCl (pH 7.4), 200 mM NaCl, and 0.1 mM PMSF. Protein concentrations were determined using the BCA protein assay. The cells expressing GFP-L2-C were lysed with Binding buffer (20mM Tris-HCl pH 7.4, 300mM NaCl) and supplemented with 20mM Imidazole, 1% Sarkosyl, 20 units/mL DNase I, 0.5 mg/mL lysozyme, and 1 mM PMSF, then purified by using a pre-equilibrated His GraviTrap column in binding buffer with 20mM Imidazole. Purified proteins were eluted from the column with binding buffer with 500 mM imidazole and exchanged into buffer containing 50 mM Tris-HCl (pH 7.4), 200 mM NaCl, and 0.1 mM PMSF by dialysis and quantified by BCA protein assay.

### GST pull-down experiments

20 μg GST or GST-Rabankyrin-5 was immobilized on glutathione (GSH) resin and incubated with 10 μg of either GFP or GFP-L2-C fusion protein in Binding buffer (20 mM HEPES, pH 7.4, 50 mM NaCl, 5 mM MgCl₂, 1 mM DTT, and 0.1% Triton X-100) at 4 °C overnight. Beads were then centrifuged and washed five times with binding buffer, resuspended in 2 × SDS loading buffer containing 2-mercaptoethanol, and heated at 95 °C for 10 minutes.

Samples were analyzed by SDS-PAGE and immunoblotting using appropriate antibodies. For the GST pulldown of dynein, the reaction was performed similarly, with 10 μg of dynein incubated with 20 μg GST or GST-Rabankyrin-5 in binding buffer containing 20 mM Tris-HCl (pH 7.4), 50 mM KCl, 2 mM MgCl₂, and 0.5 mM EGTA.

### Live cell imaging

5 × 10^3^ HeLa cells grown on eighteen-chambered glass coverslips were transfected with 10 nM negative control siRNA or siRNA targeting Rabankyrin-5. After 48 hours, transfected cells were infected with AlexaFluor488-labeled HPV.FLAG PsVs at a MOI of 100. At one hour, cells were washed three times with PBS, stained with 1 μM SiR-tubulin and 0.1 μg/mL Hoechst 33342 at 37 °C for 30 minutes, washed again with PBS, and imaged using a Nikon Ti2 spinning disk confocal microscope. Images were analyzed using the TrackMate plugin in Fiji. The mean displacement and velocity were determined using Prism by performing Gaussian fitting of the displacement frequency distributions (> 0.2 µm) and the frequency distributions of the maximum velocity of each particle with a displacement greater than 0.2 µm during a 2-minute observation period.

For imaging the co-movement of mRFP-Rab5 and Rabankyrin-5-eGFP, 1 × 10^4^ HeLa cells were plated on eighteen-chambered glass coverslips and incubated for 16 hours, then co-transfected with 0.2 μg each of mRFP-Rab5 and pcDNA3.1(+)-Rabankyrin-5-eGFP plasmids using Lipofectamine 2000, following the manufacturer’s protocol. After 48 hours, cells were infected with AlexaFluor647-labeled HPV.FLAG PsVs at an MOI of 100. At one hour, cells were stained as described above and imaged at 1.5 hpi.

### Peroxisome trafficking assay

5 × 10^4^ HeLa cells were seeded on glass coverslips in 24-well plates and co-transfected with pBa.PEX3-tdTomato-FKBP and either HA-FRB or HA-Rank-5-FRB plasmids. Twenty-four hours post-transfection, cells were treated with DMSO or 1 μM rapamycin for 1 hour. Following two PBS washes, cells were fixed with 4% paraformaldehyde for 15 minutes at room temperature. Nuclei were stained with DAPI using BD fluorescence mounting medium, and images were acquired using a Nikon Ti2 spinning disk confocal microscope. The number of cells exhibiting a clustered peroxisome phenotype near the nucleus, defined as peroxisomes relocated through microtubule-based transport activated by a dynein adaptor, was manually quantified, as shown in Fig. 5D and E.

### AlphaFold

Structural prediction of Rabankyrin-5 protein was performed using the AlphaFold Monomer v2.0 pipeline (https://alphafold.com/entry/Q9P2R3). This AI-based platform was trained on the PDB database included in AlphaFold DB version 2022-11-01. Default settings were used to generate the structure of Rabankyrin-5 protein sequence, and the model with the highest confidence score (87.1) was selected. The predicted protein structure is shown in Fig. 6A.

## ACKNOWLEDGMENTS

We thank Steve Caplan and Daniel DiMaio for reagents and helpful comments. M.L. was supported by a T32 training grant (1T32GM153375-01).

---

**Figure S1.**
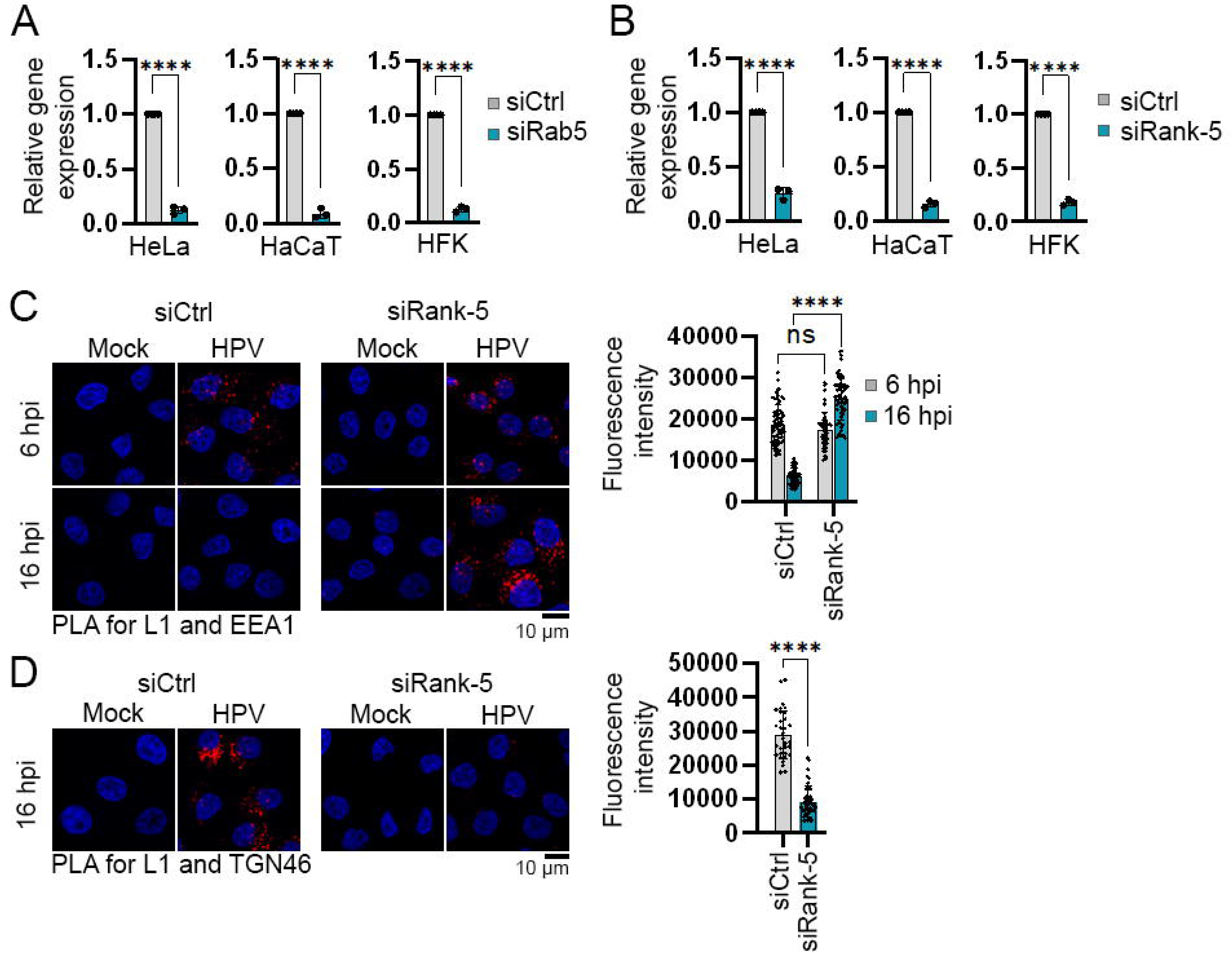
Rabankyrin-5 is required for HPV retrograde transport and is associated with HPV. (A and B) HeLa, HaCaT, and HFK cells were transfected with negative control siRNA (siCtrl), siRNA targeting Rab5A (A), or siRNA targeting Rabankyrin-5 (siRank-5) (B). Total RNA was isolated 48 hours post-transfection, and cDNA was synthesized. Expression levels of Rab5A and Rabankyrin-5 were quantified by qPCR using GAPDH as an internal control. Relative gene expression was normalized to levels observed in siCtrl-treated cells (n = 3). Statistical significance was assessed using a two-tailed, unequal variance *t*-test compared to the siCtrl group. ****p < 0.0001. (C and D) HeLa cells were transfected with siCtrl or siRNA targeting Rabankyrin-5, then infected with HPV.FLAG PsVs at an MOI of 200. At 6 and 16 hpi, PLAs were performed using antibodies recognizing HPV16 L1 and EEA1 (C) or TGN46 (D). Nuclei were stained with DAPI (blue). Scale bar: 10 μm. PLA signals (red) from at least 60 cells per condition were quantified using Fiji software. (Right) Graphs show the mean ± SD from three independent experiments; each dot represents the PLA fluorescence intensity of an individual cell. Statistical significance was determined by ANOVA. ****p < 0.0001.

**Figure S2.**
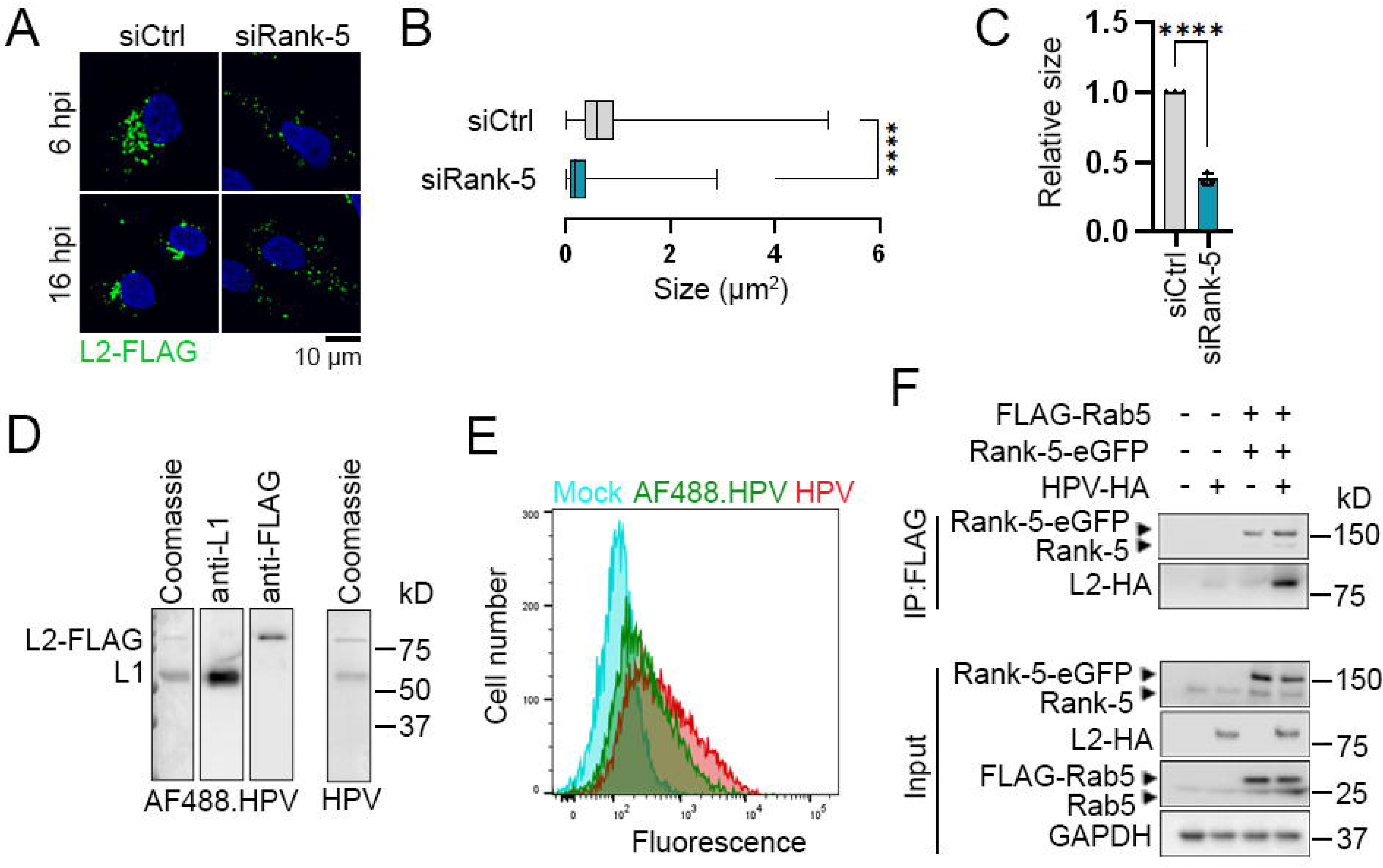
Rabankyrin-5 contributes to the fusion of HPV-carrying early endosomes. (A) HaCaT cells were transfected with either negative control siRNA (siCtrl) or siRNA targeting Rabankyrin-5 (siRank-5). After 48 hours, cells were mock-infected or infected with HPV.FLAG PsVs at an MOI of 50. Cells were stained with an anti-FLAG antibody at 6 or 16 hpi, and fluorescence was visualized by confocal microscopy. Representative images show L2-FLAG in green and nuclei in blue. Scale bar: 10 μm. (B) The size of viral puncta of infected cells at 6 hpi was quantified and plotted. Bars represent the full range of values. ****p < 0.0001; n = 1000-1200 events. (C) The relative size was determined based on the average size of virus particles shown in (B) from three independent experiments, normalized to siCtrl-treated cells (set at 1). Each dot represents the mean particle size from a single experiment. Statistical significance was assessed using a two-tailed *t*-test with unequal variance, comparing each condition to siCtrl-treated cells. ****p < 0.0001. (D) AlexaFluor 488-labeled HPV.FLAG PsVs (AF488.HPV) were analyzed by SDS-PAGE and western blotting. From left to right: Coomassie staining, western blot using an antibody against HPV16 L1, western blot using anti-FLAG antibody to detect L2-FLAG. On the right, unlabeled HPV.FLAG PsVs (HPV) were analyzed by Coomassie staining for comparison with the labeled PsVs. (E) HeLa cells were infected with either AF488.HPV PsVs or unlabeled PsVs at an MOI of approximately 2. Two days post-infection, HcRed fluorescence was measured by flow cytometry to assess infectivity. The graph shows flow cytometry histograms. (F) HeLa cells were mock-transfected (-) or transfected (+) with Rabankyrin-5-eGFP (Rank-5-eGFP) and FLAG-tagged Rab5A (FLAG-Rab5). After 48 hours, cells were mock-infected (-) or infected (+) with HA-tagged HPV16 PsVs (HPV.HA). At 6 hpi, cell lysates were immunoprecipitated using M2 FLAG beads and analyzed by western blotting for Rabankyrin-5 and HA.

**Figure S3.**
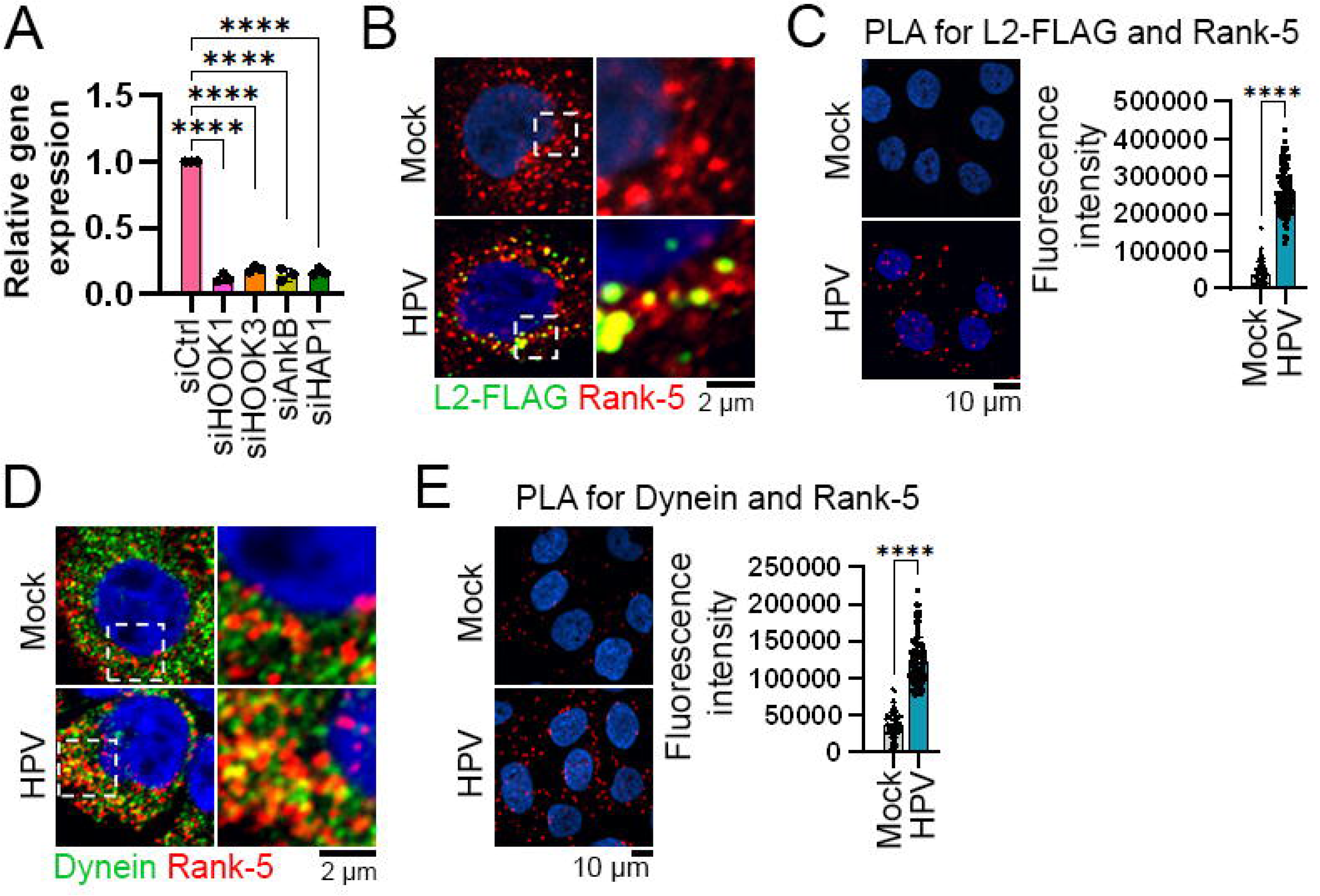
Rabankyrin-5 is in a complex with HPV L2 and dynein. (A) HeLa cells were transfected with either negative control siRNA (siCtrl) or siRNAs targeting known dynein adaptors involved in endosomal cargo transport. Total RNA was isolated 48 hours post-transfection, and cDNA was synthesized. Target gene expression levels were quantified by qPCR as described in Fig. S1A (n = 3). Statistical significance was determined using a two-tailed, unequal variance *t*-test compared to the siCtrl group. ****p < 0.0001. (B) HaCaT cells were mock-infected or infected with HPV.FLAG PsVs at an MOI of 50. At 6 hpi, cells were stained as described in Fig. 4C. Representative images (left) and insets (right) are shown. L2-FLAG (green), Rabankyrin-5 (red), and nuclei (blue) are visualized. Co-localization of L2-FLAG and Rabankyrin-5 appears yellow. Scale bar: 2 μm. (C) HaCaT cells were mock-infected or infected with HPV.FLAG PsVs at an MOI of 200. At 6 hpi, a PLA for L2-FLAG and Rabankyrin-5 was performed, as described in Fig. 4D. Each dot represents an individual cell. Statistical significance was assessed using ANOVA (****p < 0.0001). Scale bar: 10 μm. (D) HaCaT cells were mock-infected or infected with HPV16.FLAG PsVs at an MOI of 50 and stained as described in Fig. 4E. Dynein (green), Rabankyrin-5 (red), and nuclei (blue) are shown. Co-localization of dynein and Rabankyrin-5 is pseudocolored in yellow. Scale bar: 2 μm. (E) HaCaT cells were mock-infected or infected with HPV.FLAG PsVs at an MOI of 200. At 6 hpi, a PLA was performed to detect interactions between dynein and Rabankyrin-5, as described in Fig. 4F. Each dot represents an individual cell. Statistical significance was assessed using ANOVA (****p < 0.0001). Scale bar: 10 μm.

## References

1. J. Rink, E. Ghigo, Y. Kalaidzidis, M. Zerial, Rab conversion as a mechanism of progression from early to late endosomes. Cell 122, 735–749 (2005).

2. P. J. Cullen, F. Steinberg, To degrade or not to degrade: mechanisms and significance of endocytic recycling. Nat Rev Mol Cell Biol 19, 679–696 (2018).

3. Y. Yamauchi, A. Helenius, Virus entry at a glance. J Cell Sci 126, 1289–1295 (2013).

4. J. Mercer, M. Schelhaas, A. Helenius, Virus entry by endocytosis. Annu Rev Biochem 79, 803–833 (2010).

5. J. M. White, G. R. Whittaker, Fusion of Enveloped Viruses in Endosomes. Traffic 17, 593–614 (2016).

6. C. S. Kumar, D. Dey, S. Ghosh, M. Banerjee, Breach: Host Membrane Penetration and Entry by Nonenveloped Viruses. Trends Microbiol 26, 525–537 (2018).

7. M. A. Ozbun, S. K. Campos, The long and winding road: human papillomavirus entry and subcellular trafficking. Curr Opin Virol 50, 76–86 (2021).

8. J. Xie, P. Zhang, M. Crite, D. DiMaio, Papillomaviruses Go Retro. Pathogens 9 (2020).

9. Anonymous, Centers for Disease Control and Prevention. Cancers Associated with Human Papillomavirus. Centers for Disease Control and Prevention, U.S.Department of Health and Human Services; 2024. (2024).

10. P. M. Day et al., Human Papillomavirus 16 Capsids Mediate Nuclear Entry during Infection. J Virol 93 (2019).

11. S. DiGiuseppe et al., Incoming human papillomavirus type 16 genome resides in a vesicular compartment throughout mitosis. Proc Natl Acad Sci U S A 113, 6289–6294 (2016).

12. P. M. Day, C. D. Thompson, R. M. Schowalter, D. R. Lowy, J. T. Schiller, Identification of a role for the trans-Golgi network in human papillomavirus 16 pseudovirus infection. J Virol 87, 3862–3870 (2013).

13. P. Zhang, G. Monteiro da Silva, C. Deatherage, C. Burd, D. DiMaio, Cell-Penetrating Peptide Mediates Intracellular Membrane Passage of Human Papillomavirus L2 Protein to Trigger Retrograde Trafficking. Cell 174, 1465–1476 e1413 (2018).

14. J. Xie, P. Zhang, M. Crite, C. V. Lindsay, D. DiMaio, Retromer stabilizes transient membrane insertion of L2 capsid protein during retrograde entry of human papillomavirus. Sci Adv 7 (2021).

15. A. Lipovsky et al., Genome-wide siRNA screen identifies the retromer as a cellular entry factor for human papillomavirus. Proc Natl Acad Sci U S A 110, 7452–7457 (2013).

16. J. Xie, E. N. Heim, M. Crite, D. DiMaio, TBC1D5-Catalyzed Cycling of Rab7 Is Required for Retromer-Mediated Human Papillomavirus Trafficking during Virus Entry. Cell Rep 31, 107750 (2020).

17. A. Popa et al., Direct binding of retromer to human papillomavirus type 16 minor capsid protein L2 mediates endosome exit during viral infection. PLoS Pathog 11, e1004699 (2015).

18. R. Behnia, S. Munro, Organelle identity and the signposts for membrane traffic. Nature 438, 597–604 (2005).

19. S. L. Reck-Peterson, W. B. Redwine, R. D. Vale, A. P. Carter, The cytoplasmic dynein transport machinery and its many cargoes. Nat Rev Mol Cell Biol 19, 382–398 (2018).

20. A. Zeigerer et al., Rab5 is necessary for the biogenesis of the endolysosomal system in vivo. Nature 485, 465–470 (2012).

21. H. M. York et al., Deterministic early endosomal maturations emerge from a stochastic trigger-and-convert mechanism. Nat Commun 14, 4652 (2023).

22. D. H. Murray et al., An endosomal tether undergoes an entropic collapse to bring vesicles together. Nature 537, 107–111 (2016).

23. M. Schelhaas et al., Entry of human papillomavirus type 16 by actin-dependent, clathrin- and lipid raft-independent endocytosis. PLoS Pathog 8, e1002657 (2012).

24. J. L. Smith, S. K. Campos, A. Wandinger-Ness, M. A. Ozbun, Caveolin-1-dependent infectious entry of human papillomavirus type 31 in human keratinocytes proceeds to the endosomal pathway for pH-dependent uncoating. J Virol 82, 9505–9512 (2008).

25. A. Lipovsky et al., The cellular endosomal protein stannin inhibits intracellular trafficking of human papillomavirus during virus entry. J Gen Virol 98, 2821–2836 (2017).

26. C. Schnatwinkel et al., The Rab5 effector Rabankyrin-5 regulates and coordinates different endocytic mechanisms. PLoS Biol 2, E261 (2004).

27. J. Zhang et al., Rabankyrin-5 interacts with EHD1 and Vps26 to regulate endocytic trafficking and retromer function. Traffic 13, 745–757 (2012).

28. V. Nehru, O. Voytyuk, J. Lennartsson, P. Aspenstrom, RhoD binds the Rab5 effector Rabankyrin-5 and has a role in trafficking of the platelet-derived growth factor receptor. Traffic 14, 1242–1254 (2013).

29. T. Ohya et al., Reconstitution of Rab- and SNARE-dependent membrane fusion by synthetic endosomes. Nature 459, 1091–1097 (2009).

30. H. M. York et al., Rapid whole cell imaging reveals a calcium-APPL1-dynein nexus that regulates cohort trafficking of stimulated EGF receptors. Commun Biol 4, 224 (2021).

31. M. Lakadamyali, M. J. Rust, X. Zhuang, Ligands for clathrin-mediated endocytosis are differentially sorted into distinct populations of early endosomes. Cell 124, 997–1009 (2006).

32. T. A. Masters, D. A. Tumbarello, M. V. Chibalina, F. Buss, MYO6 Regulates Spatial Organization of Signaling Endosomes Driving AKT Activation and Actin Dynamics. Cell Rep 19, 2088–2101 (2017).

33. O. J. Driskell, A. Mironov, V. J. Allan, P. G. Woodman, Dynein is required for receptor sorting and the morphogenesis of early endosomes. Nat Cell Biol 9, 113–120 (2007).

34. L. Florin et al., Identification of a dynein interacting domain in the papillomavirus minor capsid protein l2. J Virol 80, 6691–6696 (2006).

35. K. Y. Lai et al., A Ran-binding protein facilitates nuclear import of human papillomavirus type 16. PLoS Pathog 17, e1009580 (2021).

36. K. Speckhart, J. Choi, D. DiMaio, B. Tsai, The BICD2 dynein cargo adaptor binds to the HPV16 L2 capsid protein and promotes HPV infection. PLoS Pathog 20, e1012289 (2024).

37. J. Miller et al., HPV16 E7 protein and hTERT proteins defective for telomere maintenance cooperate to immortalize human keratinocytes. PLoS Pathog 9, e1003284 (2013).

38. C. B. Buck, D. V. Pastrana, D. R. Lowy, J. T. Schiller, Efficient intracellular assembly of papillomaviral vectors. J Virol 78, 751–757 (2004).

39. R. Jahn, D. C. Cafiso, L. K. Tamm, Mechanisms of SNARE proteins in membrane fusion. Nat Rev Mol Cell Biol 25, 101–118 (2024).

40. M. Schelhaas et al., Human papillomavirus type 16 entry: retrograde cell surface transport along actin-rich protrusions. PLoS Pathog 4, e1000148 (2008).

41. A. Garbouchian, A. C. Montgomery, S. P. Gilbert, M. Bentley, KAP is the neuronal organelle adaptor for Kinesin-2 KIF3AB and KIF3AC. Mol Biol Cell 33, ar133 (2022).

42. W. Nijenhuis, M. M. P. van Grinsven, L. C. Kapitein, An optimized toolbox for the optogenetic control of intracellular transport. J Cell Biol 219 (2020).

43. L. C. Kapitein et al., Probing intracellular motor protein activity using an inducible cargo trafficking assay. Biophys J 99, 2143–2152 (2010).

44. R. N. Collins, J. Zimmerberg, Cell biology: A score for membrane fusion. Nature 459, 1065–1066 (2009).

45. J. A. Soler, A. Singh, M. Zerial, S. Thutupalli, Motor Function of the Two-Component EEA1-Rab5 Revealed by dcFCCS. Methods Mol Biol 2881, 87–115 (2025).

46. J. Choi, K. Speckhart, B. Tsai, D. DiMaio, Rab6a enables BICD2/dynein-mediated trafficking of human papillomavirus from the trans-Golgi network during virus entry. mBio 10.1128/mbio.02811-24, e0281124 (2024).

47. W. Zhang, T. Kazakov, A. Popa, D. DiMaio, Vesicular trafficking of incoming human papillomavirus 16 to the Golgi apparatus and endoplasmic reticulum requires gamma-secretase activity. mBio 5, e01777–01714 (2014).

